# Rapid, high-resolution, non-destructive assessments of metabolic and morphological homogeneity uniquely identify high-grade cervical precancerous lesions

**DOI:** 10.1101/2024.05.10.593564

**Authors:** Christopher M. Polleys, Pramesh Singh, Hong-Thao Thieu, Elizabeth M. Genega, Narges Jahanseir, Andrea L. Zuckerman, Francisca Rius Díaz, Abani Patra, Afshin Beheshti, Irene Georgakoudi

**Author notes:** Corresponding Author: Christopher M. Polleys, Hong-Thao Thieu, Elizabeth M. Genega, Pramesh Singh, Narges Jahanseir, Andrea L Zuckerman, Francisca Rius Díaz, Abani Patra, Afshin Beheshti, Irene Georgakoudi. Department of Obstetrics and Gynecology, Newton-Wellesley Hospital, Newton, MA 02462, USA. Department of Pathology and Laboratory Medicine, Emory University Hospital, Atlanta, GA 30322, USA.

## Abstract

**Purpose:** Two-photon microscopy (2PM) is an emerging clinical imaging modality with the potential to non-invasively assess tissue metabolism and morphology in high-resolution. This study aimed to assess the translational potential of 2PM for improved detection of high-grade cervical precancerous lesions.

**Experimental Design:** 2P images attributed to reduced nicotinamide adenine dinucleotide (phosphate) (NAD(P)H) and oxidized flavoproteins (FP) were acquired from the full epithelial thickness of freshly excised human cervical tissue biopsies (N = 62). Fifteen biopsies harbored high-grade squamous intraepithelial lesions (HSILs), 14 biopsies harbored low-grade SILs (LSILs), and 33 biopsies were benign. Quadratic discriminant analysis (QDA) leveraged morphological and metabolic functional metrics extracted from these images to predict the presence of HSILs. We performed gene set enrichment analysis (GSEA) using datasets available on the Gene Expression Omnibus (GEO) to validate the presence of metabolic reprogramming in HSILs.

**Results:** Integrating metabolic and morphological 2P-derived metrics from finely sampled, full-thickness epithelia achieved a high 90.8 ± 6.1% sensitivity and 72.3 ± 11.3% specificity of HSIL detection. Notably, sensitivity (91.4 ± 12.0%) and specificity (77.5 ± 12.6%) were maintained when utilizing metrics from only two images at 12- and 72-µm from the tissue surface. Upregulation of glycolysis, fatty acid metabolism, and oxidative phosphorylation in HSIL tissues validated the metabolic reprogramming captured by 2P biomarkers.

**Conclusion:** Label-free 2P images from as few as two epithelial depths enable rapid and robust HSIL detection through the quantitative characterization of metabolic and morphological reprogramming, underscoring the potential of this tool for clinical evaluation of cervical precancers.

**Translational Relevance Statement:** The colposcopy and biopsy paradigm for cervical pre-cancer detection leads to an excessive number of unnecessary biopsies, with significant economic and psychological costs. This study highlights the potential of label-free, high-resolution two photon imaging to improve this paradigm by introducing real-time morphofunctional tissue assessments.

In an extensive dataset comprising freshly excised high-grade and low-grade cervical intraepithelial lesions, along with benign metaplastic and inflamed human cervical tissue biopsies, we successfully characterize a loss of morphofunctional heterogeneity indicative of high-grade precancerous changes. Leveraging a combination of two-photon imaging-derived quantitative morphofunctional metrics, our findings showcase a substantial improvement in both sensitivity and specificity of high-grade lesion detection compared to the current gold standard of colposcopy followed by a single biopsy. The demonstrated enhancement in sensitivity and specificity highlights the prospect of integrating non-invasive, label-free metabolic imaging into clinical practice, offering a more effective and efficient approach to identify and manage cervical precancerous lesions.

## Introduction

Despite the success of the quadrivalent human papillomavirus (HPV) vaccine, uterine cervical cancer persists as a significant global health concern, ranking as the 4^th^ most diagnosed cancer among women worldwide^1,2^. Challenges in vaccine adoption and the imperative to safeguard individuals already infected with HPV necessitate ongoing improvements in secondary prevention methods^3,4^. In clinical practice, patients with abnormal cervical cancer screening results typically undergo colposcopy, a widely utilized procedure that involves visual examination of the cervix, and subsequent biopsy of abnormal sites. The goal of colposcopy is to locate high-grade squamous intraepithelial lesions (HSILs) for treatment, as the vast majority of LSILs either regress or persist, with a fraction of a percent progressing to invasive cancer^5,6^. Colposcopy followed by a single biopsy suffers from limited sensitivity and specificity. HSIL detection sensitivity improves from 60.6% to 85.6% and 95.6% with the acquisition of a second and third post-colposcopy biopsy, respectively^7^. Although colposcopy has the potential to achieve high sensitivity in identifying HSILs, its reliance on non-specific contrast agents poses challenges, leading to biopsies of lesions unlikely to progress to invasive cancer and of benign conditions like inflammation and metaplasia^8,9^. The use of non-specific contrast agents results in the acquisition of many unnecessary biopsies. For example, Blatt et al. indicated that HSIL+ specificity was as low as 6% in over 250,000 post-colposcopy biopsies^10^. Efforts to enhance diagnostic precision have spurred advancements in optical imaging devices for the cervix^11–13^. Nevertheless, prevailing methodologies predominantly offer superficial morphological information, leaving a substantial gap in comprehensive diagnostic capabilities.

Our study focuses on bridging this diagnostic gap by utilizing two-photon microscopy (2PM), a high-resolution, label-free imaging technique. This innovative method exploits biomolecular contrast, particularly the two-photon excited fluorescence (TPEF) generated by reduced nicotinamide adenine dinucleotide (NADH), reduced nicotinamide adenine dinucleotide phosphate (NADPH), collectively referred to as NAD(P)H, and oxidized flavoproteins (FP). Due to the differential TPEF generation efficiency of mitochondrial, protein bound NADH, NAD(P)H intensity fluctuations offer valuable insights into mitochondrial organization and overall tissue metabolic state^14,15^. As dynamic organelles responding to metabolic demands, mitochondria play a pivotal role in cellular proliferation^16^. Furthermore, the interplay between NAD(P)H and FP allows for the measurement of tissue oxido-reductive state, providing a unique avenue for understanding cancer-induced metabolic changes.

Previous work by our group demonstrated the diagnostic potential of 2P morphological and metabolic metrics in the context of the cervix, with a particular emphasis on differentiating between SIL and non-SIL tissues^17^. In this extended study, we significantly expand our dataset and refine our analysis to offer a comprehensive evaluation of HSIL detection using 2PM. We aim to develop a rapid and robust approach that optimizes the number of 2P measurements without sacrificing performance. We also aim to underscore the effectiveness of metabolic measurements, which are not currently utilized in the diagnostic scheme, especially in scenarios where false positives may arise. To validate the origins of the optical metabolic changes we detect, we supplement our findings with gene pathway enrichment analysis of publicly available cervical tissue microarray data from several teams^18–20^.

This study contributes to the evolution of cancer diagnostics by emphasizing the clinical utility of 2PM in providing both metabolic and morphological insights. By addressing the limitations of current diagnostic methods, our study presents a significant step towards improving the accuracy and efficiency of high-grade preinvasive cervical cancer detection in a clinical setting.

## Materials and Methods

### Ethical Approval and Patient Consent

All procedures pertaining to biopsy acquisition, processing, imaging, and storage were approved by the Tufts Health Sciences Institutional Review Board protocol #10283. Informed consent was obtained from all patients contributing biopsy data.

### Specimen Procurement

Premenopausal women over the age of 18 with an abnormal low-grade squamous intraepithelial lesion (LSIL) or high-grade squamous intraepithelial lesion (HSIL) Papanicolaou test undergoing colposcopy or loop electrosurgical excision procedure (LEEP) were recruited to participate in the study. Research biopsies (∼5- x 5-mm) were excised by a patient’s gynecologist using Tischler or Rongeur forceps, depending on clinician preference, from the second-most visibly abnormal region of ectocervix following the application of 3% acetic acid during the routine procedure. Premenopausal women undergoing hysterectomy for benign gynecological disease were recruited as control patients. Strips of tissue (∼5- x 25-mm) containing both endo- and ecto-cervix were sectioned from freshly excised, visibly normal uterine cervical specimens under sterile conditions by a board-certified Pathologist in the Tufts Medical Center (TMC) department of Pathology and Laboratory Medicine. Tissue samples were transported back to the Tufts Advanced Microscopic Imaging Center (TAMIC) in a specimen cup containing a custom-built tissue carrier and a small volume of keratinocyte serum-free medium (Lonza) to provide protection and physiologically relevant hydration. Samples were imaged within 4-hours of excision. Biopsies were placed in 10% neutral buffered formalin following imaging and were returned to the TMC department of Pathology and Laboratory Medicine for standard histopathological diagnosis. For five biopsies, tattoo inks were used to mark the epithelial surface, allowing for the determination of TPEF imaging locations. One H&E section was acquired per optical region of interest (ROI). In such cases, all ROIs from a single biopsy had an agreement in diagnosis (Supplemental Methods, Supplemental Figure S1). For all other biopsies, a diagnosis for all optical ROIs was rendered from one hematoxylin and eosin (H&E)-stained tissue section.

### Patient Cohort

Eighty-eight (88) total patients consented to participate in the study between 2019 and 2023. Twenty-six (26) patients were excluded due to issues of quality control and specimen access (Supplemental Methods, Supplemental Table S1). Data from 62 patients were included for analysis (Table 1). All 19 samples resected from hysterectomy specimens contained benign squamous mucosa. Of the LEEP and colposcopy biopsies, 15 were HSIL, 14 were LSIL, and 14 were benign.

**Table 1.** Patient demographic information.

| **Characteristic** |  | **Cohort**<br>**N = 62** |
| --- | --- | --- |
| Age | 21 – 30 | 23 (37.1%) |
|  | 31 – 40 | 22 (35.5%) |
|  | 41 – 50 | 16 (25.8%) |
|  | 51+ | 1 (1.6%) |
| Race | Asian | 13 (21.0%) |
|  | Black or African American | 11 (17.7%) |
|  | White | 33 (53.2%) |
|  | Unknown | 5 (8.1%) |
| Diagnosis | Benign | 33 (53.2%) |
|  | LSIL | 14 (22.6) |
|  | HSIL | 15 (24.2) |
| Metaplasia |  | 10 (16.1%) |
| Inflammation |  | 14 (22.6%) |
| HPV-Type | 16/18/45 | 17 (27.4%) |
|  | 31/33/35/39/51/52/56/58/59/66/68 | 18 (29.0%) |
|  | None/Not Reported | 27 (43.6%) |

### Imaging

A commercial Leica SP8 microscope equipped with a femtosecond laser (Insight, Spectra-Physics) was used for imaging. The microscope utilized an inverted scheme in which light was delivered and collected with a 40X/1.1 numerical aperture (NA) water-immersion objective lens. Twelve-bit depth images were formed using bidirectional raster scanning at a rate of 600 lines per second. At each depth within the tissue, six individual 1024- x 1024-pixel (290- x 290-μm) image frames were acquired. Images were sampled every 4-μm through the full thickness of the tissue epithelium. Tissue volumes were first excited in their entirety with 755 or 775 (755/775) nm light, and then subsequently with 860 nm light. 16 specimens were excited with 775 nm light to compare metabolic readouts with those acquired with 755 nm, the NAD(P)H excitation wavelength our group traditionally uses. Studies by other groups have indicated that NAD(P)H fluorescence signatures do not vary over this excitation range^21,22^. Incident laser power was linearly varied through depth, with approximately 10 mW being delivered to superficial optical sections and 60 mW to basal cell layers. A previous study determined that 60 mW of near-infrared light, focused with a 40X/1.3 NA oil-immersion objective lens, from a femtosecond laser delivers a 0.6 minimal erythema dose (MED), equivalent to approximately 15 minutes of summer sun-exposure; by comparison, 1.0 MED is the threshold of sun burn development^23^. Thus, an incident power threshold of 10 – 60 mW allowed for the acquisition of high signal-to-noise ratio images at safe irradiation levels. Two to four tissue volumes, or ROIs, were imaged per biopsy. Images were collected with two hybrid photodetectors (HyDs) and two photomultiplier tubes (PMTs). One HyD was equipped with a 460/50 nm bandpass filter and the other with a 525/50 nm bandpass filter. One PMT was equipped with a 430/20 nm bandpass filter and the other with a 624/40 nm bandpass filter.

### Image Processing

Signal acquired from the 460/50 nm bandpass-filtered HyD during 755/775 nm excitation was attributed to NAD(P)H two-photon excited fluorescence (TPEF)^17^. Signal acquired from the 525/50 nm bandpass-filtered HyD during 860 nm excitation was attributed to oxidized flavoprotein (FP) TPEF^24^. Signal acquired from the 624/40 nm bandpass-filtered PMT during 860 nm excitation was attributed to hyperfluorescent cells that were removed from analysis. Signal acquired from the 430/20 nm bandpass-filtered PMT during 860 nm excitation was attributed to collagen second harmonic generation (SHG). SHG is a 2^nd^ order non-linear scattering process produced by non-centrosymmetric molecules, such as collagen, where two incident photons are simultaneously upconverted into a single photon of exactly twice the energy. All raw pixel intensities were normalized by detector gain and squared laser power. Each of the six individual frames acquired at a particular depth were averaged. Images were down-sampled into 512- x 512-pixels. All 755/775 nm excitation and 860 nm excitation images were co-registered in 3D using data collected from the 525/50 nm bandpass-filtered HyD (Supplemental Methods, Supplemental Figure S2).

Several masks were generated automatically to isolate cellular cytoplasm (Supplemental Figure S3). The intraepithelial tissue region was defined based on percent cell coverage and signal-to-noise ratio (SNR) (Supplemental Figure S4). SNR was calculated as 10 times the log base 10 of the ratio between the mean NAD(P)H power spectral density (PSD) for frequencies corresponding to 7 – 50-μm length scales, the approximate size of cells and, thus, attributed to the signal of interest, and the NAD(P)H PSD for the highest discrete spatial frequency, corresponding to a measure of noise. A PSD curve was generated by taking the squared amplitude of the 2D Fourier transform of an image, and it quantified the relative contributions of discrete spatial frequencies.

Morphological metrics of epithelial thickness and differentiation gradient were calculated on a per ROI basis. Epithelial thickness was defined as the distance from the superficial cell layer to the depth at which epithelial cells occupied more than 30% of the field of view (as opposed to collagen and stromal cells). The differentiation gradient was calculated from integrated TPEF intensity images, defined as the sum of the FP and NAD(P)H image channels. The log base 10 of the noise-normalized PSD was calculated for each integrated TPEF image in a ROI. Nuclear and cell borders feature prominently in these PSDs (Supplemental Figure S5). The variance of the noise-normalized PSD at each discrete spatial frequency was calculated for either all included depths, or the absolute depths specified. The coefficient of variation of the PSD variance across the epithelial depth for frequencies corresponding to 7 – 50-μm length scales (i.e. the typical size of cells from the basal to the superficial layer) was reported as our metric of differentiation gradient. The PSD generated by a TPEF image in this manner was sensitive to changes in the length scales of morphological features such as nuclei and cell size. A high differentiation gradient corresponded to high morphological variability between cell layers^17,25,26^.

The cellular cytoplasm mask was used to extract metabolic tissue metrics of optical redox ratio (RR) and mitochondrial clustering. Metrics of RR were extracted from cytoplasm-positive regions using the pixel-wise intensity relationship outlined in equation 1.

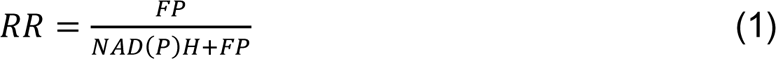

The mean and interquartile range (IQR) of the RR distribution for a given optical section were used as measures of overall and intrafield heterogeneity of tissue oxido-reductive state, respectively.

Mitochondrial clustering was calculated from cytoplasm-positive variations in NAD(P)H intensity, as previously described^15,25–28^ (Supplemental Figure S6A). Although NAD(P)H can be either mitochondrial or cytosolic, NAD(P)H bound to enzymes of the Kreb’s cycle and the electron transport chain (ETC) fluoresces 2 – 10-times more efficiently than free NAD(P)H^14^. For this reason, bright NAD(P)H fluorescence is assumed to emanate primarily from mitochondria, with NAD(P)H intensity fluctuations informing mitochondrial organization. To provide a quantitative metric of this organization relying on a fast, robust, and relatively simple to implement approach, it is important to remove from the images prominent features associated with cell and nuclear borders. For this reason, following identification of cell cytoplasmic regions, we populated void regions (Supplemental Figure S6B) by randomly cloning cytoplasm-positive pixels across the field of view until full (Supplemental Figure S6C). The Fourier-analysis based PSD was utilized once more to quantify the relative abundance of features corresponding to different characteristic sizes or spatial frequencies. The average PSD from five cloned images was fit with equation 2 for frequencies (k) corresponding to length scales less than 8.5-μm.

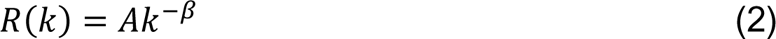

The absolute value of the fitted exponent (β) was reported as a quantitative metric of mitochondrial clustering, as several previous studies performed with cell monocultures and living tissues have shown that higher β values correspond to higher levels of mitochondrial fragmentation^15,25,26^ (Supplemental Figure S6D). The mean, median, variance, and range were calculated for metrics of RR, RR IQR and mitochondrial clustering for either all included depths, or the absolute depths specified. All image processing steps were completed using MATLAB version 2021b.

### Statistical Analysis and Classification

Statistical comparisons were made between HSIL and non-HSIL tissues for all metabolic and morphological metrics derived from the full epithelial thickness. Comparisons were also made between mature and immature non-HSIL tissues. Well-differentiated, mature benign tissues contained superficial exfoliating layers characterized by low SNR in the NAD(P)H TPEF channel. Benign tissues were classified as mature if the first high SNR (>9) optical section was beyond 60-μm into the tissue. All statistical comparisons were made in SAS JMP Pro 16. Nested t-tests were used to make statistical comparisons at the patient level while considering the intratissue variations among multiple ROIs.

The MATLAB Classification Learner application, which includes decision trees, discriminant analyses, support vector machines, low-level machine learning algorithms, and several other classifiers, was used to select the optimal diagnostic framework. Non-colinear metrics (r < 0.7) with the highest statistically significant differences between HSIL and non-HSIL tissues were used as predictor variables. An 80/20 train-test split, and 5-fold cross validation were used to evaluate classifiers. An 80/20 train-test split was used to provide sufficient training data for the machine learning-type models. Exploratory quadratic discriminant analyses (QDAs), which utilized no train-test split, were used to determine final predictor variables. Exploratory QDAs considering full thickness tissue metrics were evaluated in SAS JMP Pro 16. Exploratory QDAs considering metrics derived from 2-, 3-, and 4-optical section combinations were evaluated in MATLAB. Performance for 2-, 3-, and 4-depth exploratory QDAs was assessed using the sum of HSIL sensitivity and specificity. The highest performing 2-depth combinations that included all or 80% of the total specimens were further investigated. Featured 3- and 4-depth combinations were the best performing combinations that included the previous depths. ROIs were held constant for alike 2-, 3-, and 4-depth combinations. Predictive QDAs, which leveraged 10 randomly initialized seeds with equal class-proportioned 70/30 train/test splits, were evaluated in MATLAB. Whether classification was done in SAS JMP Pro 16 or MATLAB was based on ease of use. MATLAB facilitated the ability to rapidly iterate through multiple randomly initialized seeds and multiple depth combinations. A 70/30 train-test split was used to provide a sufficient test set for predictive QDAs.

### Gene expression analysis

Differentially expressed genes across HSIL and non-HSIL tissues were identified using the web-based tool *GEO2R* with default settings separately for three different gene expression data sets of cervical epithelial samples. In the first data set (obtained from Gene Expression Omnibus (GEO) using accession number GSE27678 and Platform GPL571), the HSIL group contained 21 HSIL samples and the non-HSIL group contained 12 benign and 11 LSIL samples. For the second data set (obtained from GEO using accession number GSE63514), the HSIL group was formed by 22 cervical intraepithelial neoplasia grade 2 (CIN2) and 40 CIN3 samples, whereas the non-HSIL group contained 24 benign and 14 CIN1 samples. For the third data set (obtained from GEO using accession number GSE7803), the HSIL group contained 7 HSIL samples and 10 benign samples). For all data sets, the differential expression of genes in the HSIL group as compared to the non-HSIL group was measured in terms of *t-score*.

### Pathway enrichment

The gene *t-scores* were used to rank the differentially expressed genes. The ranked list of genes was subsequently used to perform pathway enrichment for each gene expression data set. Pathway enrichment was performed using R-package *fGSEA*^29^ against two different pathway gene sets: (i) hallmark gene sets downloaded from the Human Molecular Signatures Database (MSigDB)^30^ and (ii) custom mitochondrial pathway gene sets^31^. Using *fGSEA*, a Normalized Enrichment Score (NES) was obtained for each pathway and its statistical significance was determined by Benjamini-Hochberg (BH)-adjusted p-value (*padj*), also called False Discovery Rate (FDR). Results were reported for pathways with *padj* ≤ 0.25.

## Results

### Label-free TPEF metrics capture metabolic and morphological perturbations in HSIL tissues

Redox ratio-coded images from representative benign, LSIL, and HSIL tissues at several depths, and their corresponding depth trends in metabolism are illustrated in Fig. 1A. Cells in 2P metabolic images are characterized by dark nuclei and bright cytoplasm. Cells at 20- and 60-μm in benign and LSIL tissues are visibly larger than those in HSIL tissues. At a depth of 120-μm, cell size is largely similar between the three groups. It can also be noted that the hues in the images, which have metabolic implications, are more uniformly blue-shifted, indicating a lower redox state, as a function of precancer status.

**Figure 1.**
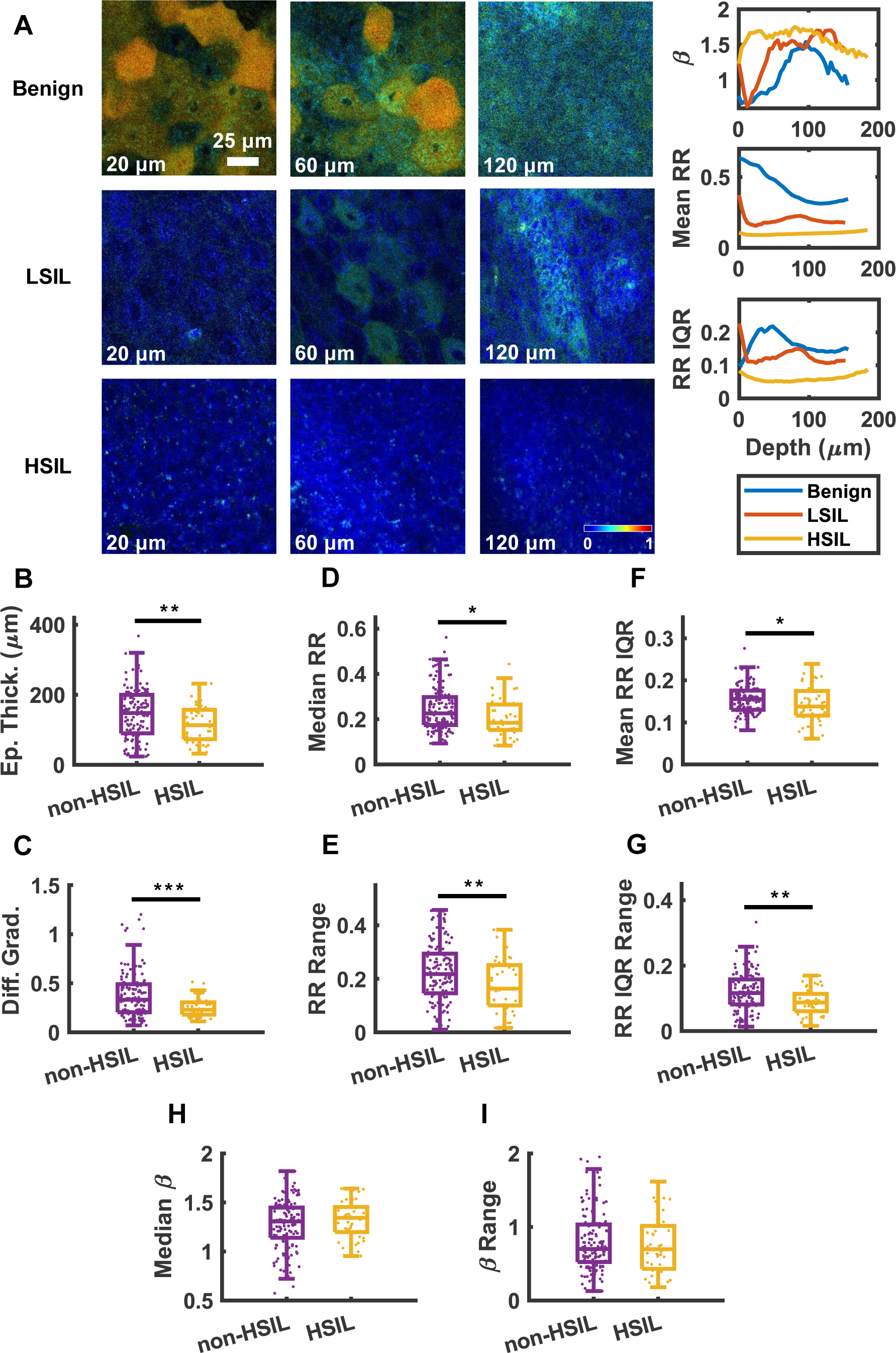
HSIL tissues differ from non-HSIL tissues in metabolism and morphology. (A) Representative redox ratio-coded images and the corresponding traces of mean redox ratio, redox ratio IQR, and mitochondrial clustering as a function of depth for benign, LSIL, and HSIL tissue biopsies. Morphologically, HSIL tissues are thinner (B) and lack depth-dependent variation in N:C (C). In terms of metabolic function, HSIL tissues are characterized by overall lower levels of oxidation (D). HSIL tissues are also characterized by a lack of heterogeneity in redox state both laterally (F) and as a function of depth (E, G). Although not significantly different, HSIL tissues are characterized as having more fragmented mitochondria (H), more homogenously distributed through the epithelium (I).

Metrics pertaining to tissue morphology and function were extracted from the full collection of intraepithelial TPEF images acquired from a single region of interest. Although it is not a traditional morphological indicator of cervical intraepithelial neoplasia, HSIL tissues exhibited a thinner epithelium compared to non-HSIL tissues (Fig. 1B). This observation can be attributed to the many exfoliating cell layers present in areas of fully differentiated benign tissue. Precancerous lesions of the human cervix are traditionally diagnosed using depth-dependent changes in intraepithelial nuclear-to-cytoplasm (N:C) ratio as visualized by hematoxylin and eosin-stained tissue cross-sections. Reduced variation in N:C as a function of depth in HSIL tissues was captured by the TPEF-based metric of differentiation gradient (Fig. 1C). Thus, morphological metrics derived from TPEF images capture known high-grade precancerous change and can do so in a non-invasive, label-free manner.

The use of NAD(P)H and FP as endogenous sources of contrast also allows for the measurement of tissue metabolic state for HSIL vs. non-HSIL tissues. The involvement of NAD(P)H and FP in several metabolic pathways including, but not limited to, glycolysis, fatty acid oxidation, glutaminolysis, and oxidative phosphorylation allow for functional conclusions to be derived from TPEF images. Functional metabolic metrics such as mitochondrial clustering, mean RR, and RR IQR (a metric of RR heterogeneity) were extracted on a per image basis. Summary metrics such as the mean, median, sample variance, and range were derived from the full collection of images as a function of depth for 3D tissue volumes (Fig. 1D - I).

All summary metrics associated with RR carried statistically significant differences between HSIL and non-HSIL tissues. In terms of RR variations, the RR range, RR IQR range, and mean RR IQR values were significantly lower in HSIL tissues compared to non-HSIL tissues. RR Range represents a measure of RR heterogeneity as a function of depth. Mean RR IQR represents an absolute measure of lateral heterogeneity present in the tissue. RR IQR range measures the variation of lateral heterogeneity as a function of depth. Our results indicate that cell layers in HSIL tissues are more homogenously aligned in metabolic state, both laterally and as a function of depth, due to the occupation of proliferative cells spanning the full epithelial thickness (Fig. 1E - G). Median redox ratio was significantly lower in HSIL tissues compared to non-HSIL tissues. A decrease in redox ratio can be attributed to several metabolic perturbations including hypoxia, enhanced fatty acid metabolism, and activation of glycolytic pathways^32^ (Fig. 1D). Metrics of median redox ratio and mean redox ratio IQR were colinear, and not simultaneously included in classification schemes that are described later. Metrics of mitochondrial clustering did not carry statistically significant differences between HSIL and non-HSIL tissues. However, HSIL tissues trended towards higher (p = 0.14, Fig. 1H) and less varied (p = 0.37, Fig. 1I) values of mitochondrial clustering through depth. Such trends in mitochondrial clustering indicate a more homogenous distribution of fragmented mitochondria (associated with enhanced glycolysis) spanning the full thickness of HSIL tissues. Together, these results highlight that functional metrics derived from TPEF images are consistent with an overall increase in the activity of glycolysis and/or fatty acid oxidation compared to oxidative phosphorylation within high-grade precancerous changes leading to significant decreases in metabolic heterogeneity present throughout the epithelium when compared to non-HSIL tissues.

### A loss of metabolic and morphological heterogeneity is essential to differentiating HSILs from LSILs and less-differentiated benign tissues

Comparisons between “mature” and “immature” benign epithelia were drawn to highlight the relevance of the present non-HSIL dataset to benign lesions that are most frequently biopsied because their morphology bears similarities to HSILs^8,9^. Specifically, we aimed to emphasize the importance of making comparative non-HSIL measurements from locations within the transformation zone, as opposed to nearby tissue, further from the cervical OS. Benign tissue stacks characterized by low N:C superficial cells with pyknotic nuclei and an exfoliating region exceeding 60-μm were classified as mature, more-differentiated benign tissues. Included in the immature class were benign tissues containing squamous metaplasia and non-metaplastic tissues with higher N:C superficial cells. Redox ratio-coded images from representative mature and immature benign tissues at several depths, and their corresponding depth trends in metabolism are illustrated in Fig. 2A. Cells at 20- and 60-μm are not as large in less-differentiated benign tissues. The image hues are also more uniformly blue-shifted. Observing the metabolic metrics’ dependence on depth, we note that immature benign tissues followed trends similar to LSIL tissues. The differences in tissue morphology that motivated the distinction between mature and immature benign tissues were reflected in the morphological TPEF metrics. Mature benign tissues were significantly thicker than immature benign tissues, supporting the claim that differences in epithelial thickness between HSIL and non-HSIL tissues can be attributed to the exfoliating regions of more differentiated benign regions (Fig. 2B). The differentiation gradient of mature benign tissues was significantly greater than that of immature benign tissues. This observation is consistent with the expectation that more differentiated benign tissues would have a greater range of N:C ratios through depth (Fig. 2C). Mature and immature benign tissue stacks also demonstrated differences in terms of extracted metabolic function metrics. The trends in mitochondrial clustering, mean RR, and RR IQR for immature benign tissues as a function of depth presented similarly to those of LSIL and HSIL tissues. The median and range of redox ratio values were significantly lower in immature benign tissues compared to mature benign tissues. Immature benign tissues are not fully differentiated and therefore contain proliferative cells spanning the full thickness of the epithelium. As expected, immature tissues exhibited a lower range and absolute level of RR values through depth, more consistent with SIL tissues (Fig. 2D-E). After removing the 23 mature non-HSIL tissues stacks, only metrics of differentiation gradient, redox ratio range, and redox ratio IQR range remained significantly different between HSIL and non-HSIL tissues (Fig. 2F). Thus, despite the similarities between HSIL and immature non-HSIL tissues, metrics of tissue differentiation and RR spatial heterogeneity persist as the major biomarkers of high-grade cervical precancerous changes.

**Figure 2.**
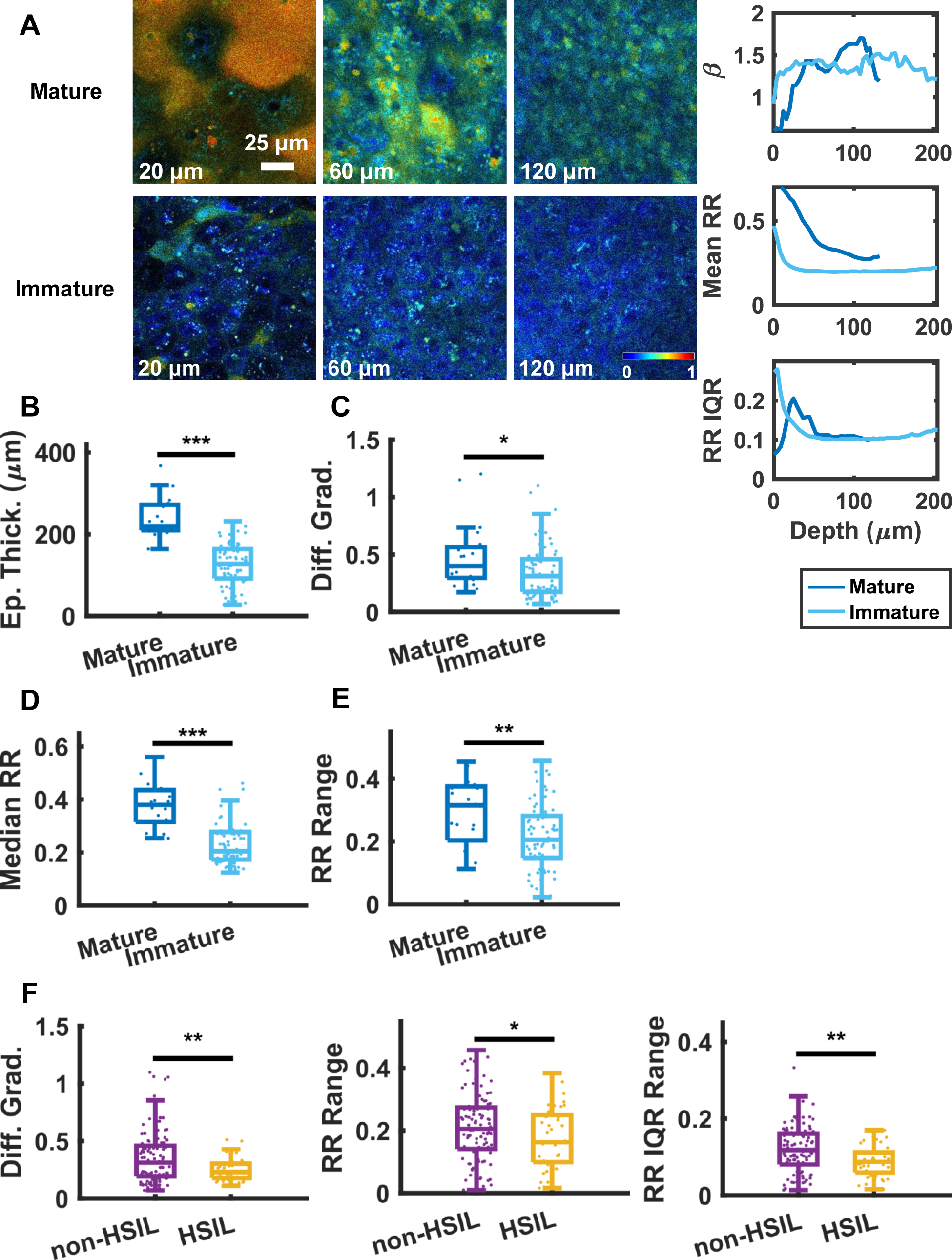
Diagnostically challenging, immature benign tissues are markedly different from well-differentiated, mature benign tissues; HSIL tissues and immature non-HSIL tissues differ in metabolism and morphology. (A) Representative redox ratio-coded images and the corresponding traces of mean redox ratio, redox ratio IQR, and mitochondrial clustering as a function of depth for mature and immature benign tissue biopsies. Morphologically, immature non-HSIL tissues are thinner and have a lower variation in N:C (C) compared to mature non-HSIL tissues. In terms of metabolic function, immature non-HSIL tissues are characterized by overall lower (D), more homogenous (E) levels of oxidation. HSIL tissues are still characterized by lack of variation in N:C and lack of metabolic heterogeneity as a function of depth (F).

### Quantitative, label-free morphofunctional metrics enable HSIL detection with high sensitivity and specificity

MATLAB Classification Learner models were tested both with and without the inclusion of mitochondrial clustering metrics. The quadratic discriminant analysis (QDA)-based classifier that included metrics of mitochondrial clustering yielded the highest validation accuracy, and therefore motivated the use of QDA for future classification investigations. In the exploratory QDAs, addition of each morphofunctional metric improved the receiver operating characteristic (ROC) area under the curve (AUC) (Table S2). Ultimately, based on stepwise variable selection, the metrics considered for predictive QDA were epithelial thickness, differentiation gradient, median RR, RR Variability, RR IQR Range, β variability, and median β. When discriminating from mature-containing non-HSIL tissues, HSILs were identified with a 90.8 ± 6.1% sensitivity and 72.3 ± 11.3% specificity (Fig. 3A). Thus, HSIL sensitivity achieved using label-free, TPEF imaging-based metrics was comparable to the sensitivity achieved by the acquisition of multiple invasive biopsies^7^, while specificity was significantly improved compared to the wide range of specificities reported for colposcopy and biopsy (6% - 88%)^8,10,33^. However, this specificity comparison is not necessarily a direct one, since for the colposcopy and biopsy studies, most of the non-HSIL biopsied tissues were immature epithelia. When we removed fully differentiated mature epithelia from our non-HSIL group, HSIL sensitivity (83.8 ± 9.2%) and specificity (69.7 ± 9.0%) were maintained. Similarly, the introduction of each morphofunctional metric improved ROC AUC (Table S3). Also of interest, metric diagnostic importance order was slightly modified (Fig. 3B). The RR IQR Range became the most important factor, highlighting the loss of metabolic heterogeneity, as opposed to morphological heterogeneity, as the principal factor in the identification of HSILs in this case. Collectively, these results demonstrate that the combination of quantitative tissue metabolic dysfunction metrics with tissue morphology characteristics yields high sensitivity and specificity of HSIL detection, even when compared.

**Figure 3.**
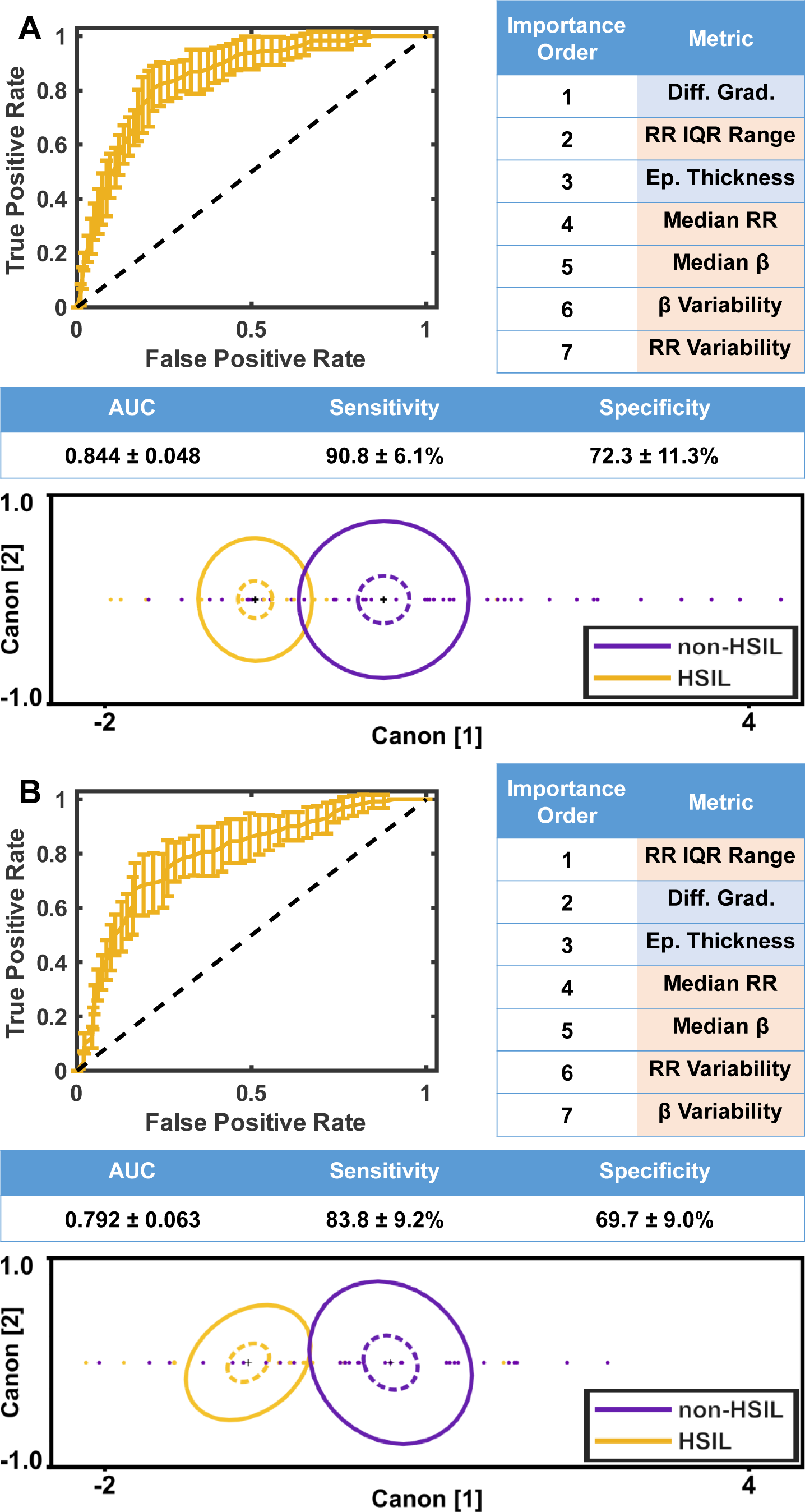
A combination of metabolic and morphological 2P biomarkers derived from the full thickness epithelium classify HSILs with high sensitivity and specificity. Receiver operating characteristic (ROC) curves are generated from quadratic discriminant analysis-based classifiers of HSILs vs. all non-HSILs (A) and immature non-HSILs (B). The importance order of metabolic and morphological metrics, ROC area-under-the-curve, sensitivity, and specificity slightly change with the exclusion of mature, benign tissues. Metabolic and morphological metrics are highlighted in orange and blue, respectively. Representative canonical plots from one randomly initialized seed are plotted, where each point represents an ROI in the test dataset. Dotted lines represent 95% confidence intervals and solid lines contain 50% of the population for each group.

### Highly accurate HSIL detection is maintained with morphofunctional assessments from two depth-resolved, label-free TPEF images

When considering clinical translation of 2PM, measurements must be fast and accurate. For this reason, it is important to identify the optimal number of optical sections that can be sampled without compromising diagnostic performance, enabling measurements within a few seconds. For models that considered data from all 15 HSIL specimens, using metrics from a combination of 3 depths had the highest AUC (0.683 ± 0.039), and therefore may the be most diagnostically useful (Fig. 4A - B). Considering depth combinations inclusive of all HSIL tissues was limiting, as some lesions only extended as far as 40-µm. The average epithelial thickness for HSIL tissues was 110 ± 50-µm. We aimed to evaluate a depth combination in this range, to be more representative of the full dataset. The 2-depth combination of 12- and 72-µm had a high combined sensitivity and specificity during exploratory QDA and considered 80% of the specimens, so this depth combination was investigated further. HSIL sensitivity (91.4 ± 12.0%) and specificity (77.5 ± 12.6%) using measurements from 12- and 72-µm outperformed the similar 3- and 4-depth combinations. In this dataset, additional diagnostic accuracy was not afforded through the acquisition of additional measurements, motivating the use of just 2 depths during clinical implementation (Fig. A - B). Models were also evaluated without the use of mitochondrial clustering metrics, as their extraction relies in principle on high resolution imaging (Fig. 4C). For models considering measurements from 12- and 72-μm, the use of mitochondrial clustering metrics improved the ROC AUC by 0.08 (Fig. 4). These results suggest that a two-depth sampling scheme from 1 superficial and 1 deep optical section is suitable for HSIL detection. A clinical imaging device should prioritize high NA acquisition, with the aim of measuring the diagnostically useful metric of mitochondrial clustering.

**Figure 4.**
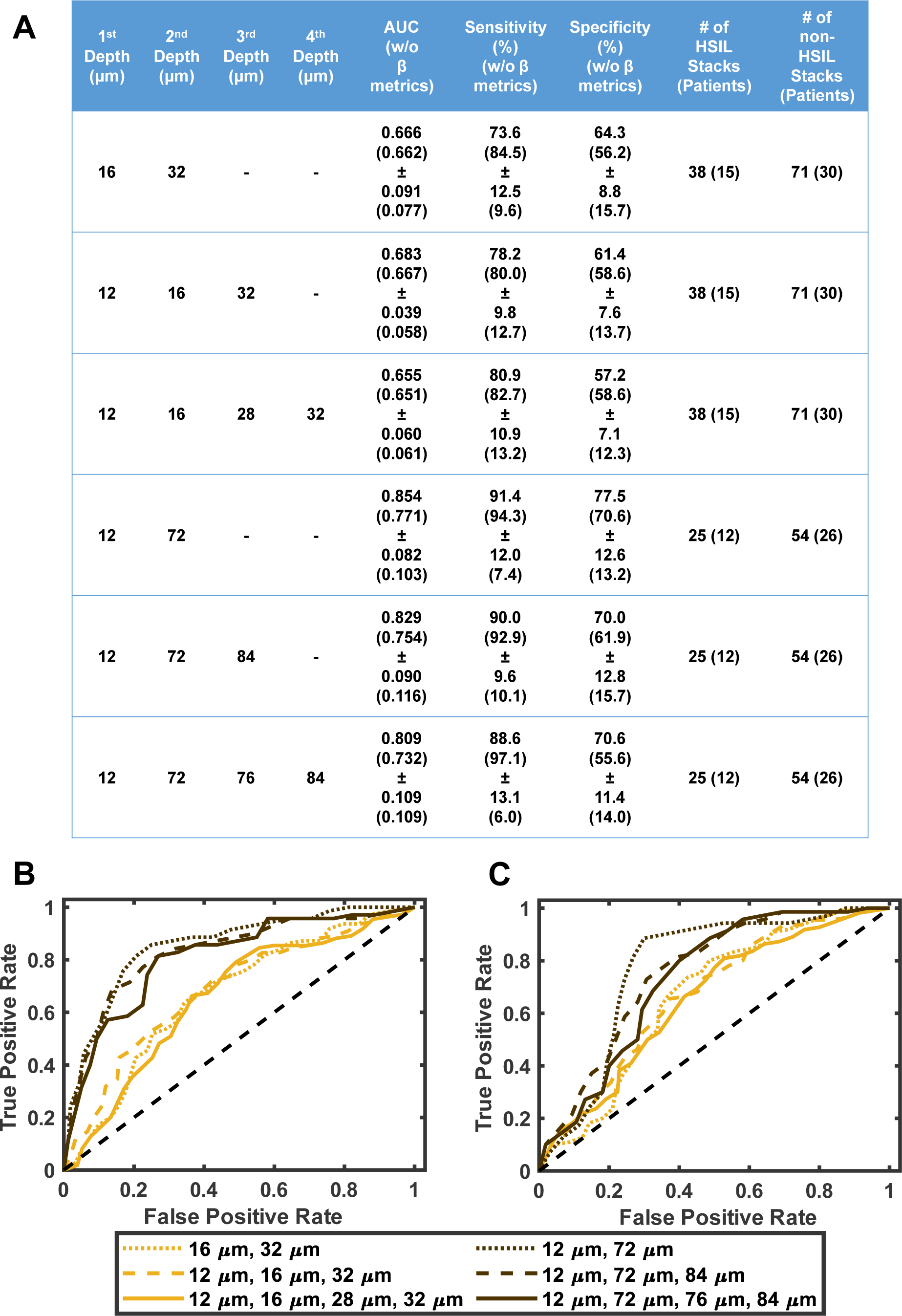
High sensitivity and specificity HSIL detection is achieved after reducing the number of depth-resolved optical sections considered for metric extraction. (A) Receiver operating characteristic (ROC) curves are generated from quadratic discriminant analysis-based classifiers of HSILs vs. immature non-HSILs when considering metrics derived from the listed depths. (B) The performance of algorithms considering morphological, redox ratio and mitochondrial organization-based metrics was similar for combinations derived using images of sections at two, three or four different depths. Utilization of information from a combination of shallow (12-μm) and deeper (72-μm) resulted in optimal classification. (C) The performance of algorithms considering only morphological and redox ratio-based metrics was reduced by very moderate levels compared to the algorithms in panel (B) for combinations derived using images of sections at two, three or four different depths. Utilization of information from a combination of shallow (12-μm) and deeper (72-μm) resulted in optimal classification.

### Pathway analysis captures HSIL metabolic complexity and validates metabolic image readouts

Custom gene sets^31^ were used to assess differential gene expression related to mitochondrial metabolism in HSIL tissues from three independent datasets, GSE27678^19^, GSE63514^20^, and GSE7803^18^. Oxidative phosphorylation (OXPHOS) activity was enhanced for HSILs in all three datasets (Fig. 5A). Additionally, glycolysis and fatty acid oxidation (FAO) were upregulated for HSIL tissues in two and one dataset, respectively (Fig. 5A). Anabolic pathways, including hypoxia inducible factor (HIF)^34,35^, mammalian target of rapamycin (mTOR)^36^, and nucleotide synthesis were also upregulated for HSIL tissues in all three datasets (Fig. 5A). Upregulation of the peroxisome and antioxidant defense pathways indicate the presence of reactive oxygen species (ROS) for HSILs in two of the datasets (Fig. 5A). Moreover, cellular stress pathways such as the unfolded protein response (UPR) and mitophagy were upregulated for HSIL tissues in three and one dataset, respectively (Fig. 5A).

**Figure 5.**
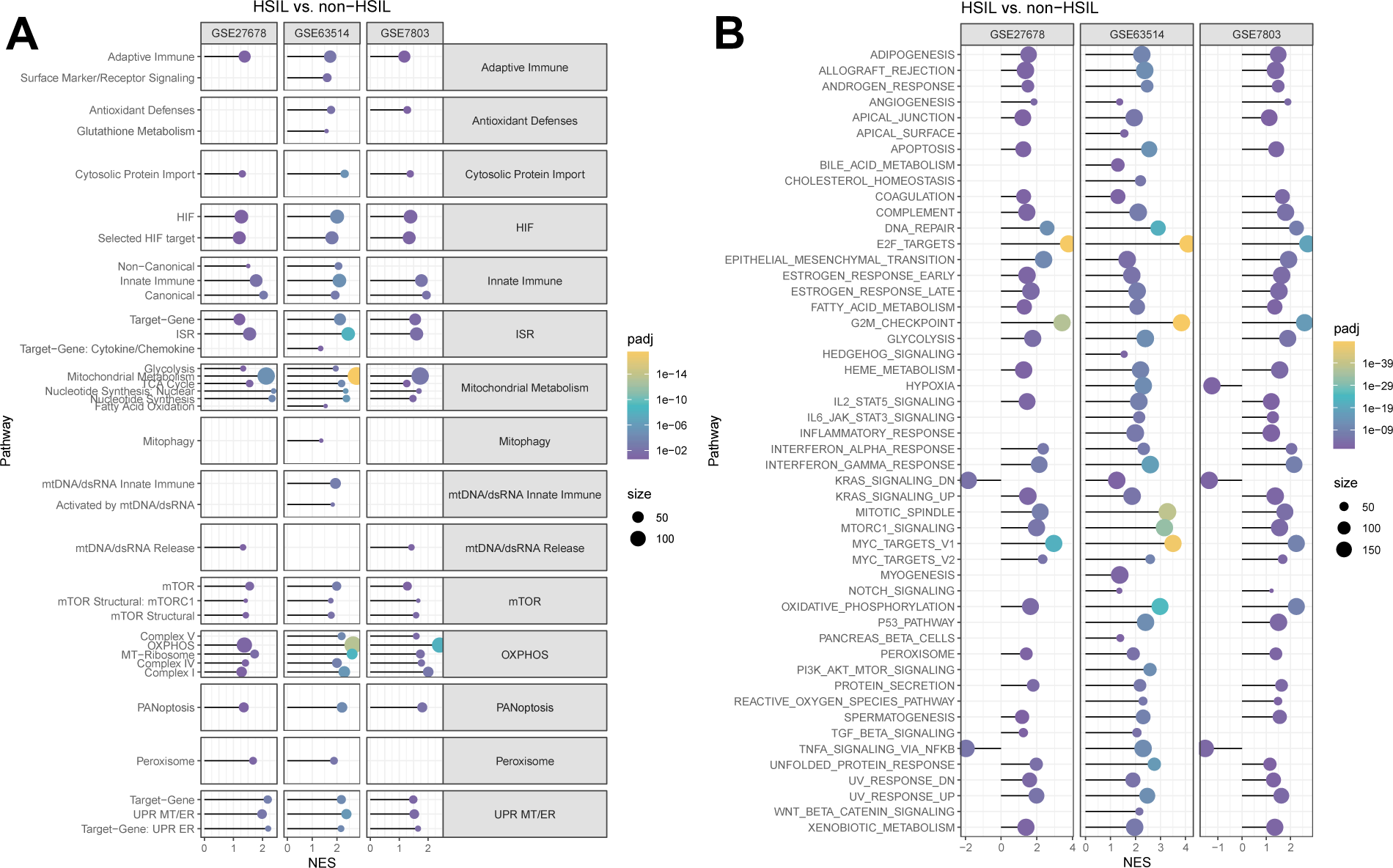
Pathway enrichment in HSIL tissues. Fast gene set enrichment analysis (fGSEA) using custom mitochondrial (A) and Hallmark (B) gene sets was used to compare HSIL and non-HSIL tissues in 3 independent datasets (GSE27678, GSE63514, and GSE7803). A positive normalized enrichment score (NES) indicates that a particular pathway is upregulated in HSIL tissues compared to non-HSIL tissues. Dot size corresponds to the size of the corresponding gene set. The dot pseudocolor is scaled based on the calculated Benjamini-Hochberg (BH)-adjusted p-value.

MSigDB Hallmark gene sets broadened the assessment of metabolic pathway analysis^30^. Consistency in the differential expression of OXPHOS, mTOR and UPR pathways was observed between Hallmark and custom mitochondrial gene sets (Fig. 5B). Hallmark gene sets indicated increased activity of glycolysis and fatty acid metabolism in all datasets (Fig. 5B). Unique to Hallmark, Myc, transcription factor E2 targets, and G2/M cell division checkpoint pathways, additional indicators of increased cellular proliferation, were upregulated in all datasets^30,37,38^ (Fig. 5B). Conflicting findings emerged for the Hallmark hypoxia pathway, showing upregulation in one dataset, downregulation in another, and no change in the third (Fig. 5B). The Hallmark ROS and peroxisome pathways were upregulated in two and three datasets, respectively, aligning with increased antioxidant defense from custom sets (Fig. 5B).

In summary, fGSEA indicates that HSIL tissues have significant anabolic demands, leverage several pathways for catabolic ATP generation, and experience several forms of cellular stress.

## Discussion

Label-free, 2PM is emerging as a transformative clinical imaging technique, with the potential to offer insights into single cell metabolism and morphology in living tissues non-invasively. 2PM has been extensively studied *in vivo* for dermatological conditions such as melanoma, basal cell carcinoma, melasma, and vitiligo^26,39,40^. This study unveils the translational potential for the rapid and accurate diagnosis of HSILs using signals derived from NAD(P)H and FP TPEF, supported by a robust dataset of 15 HSIL patients, 14 LSIL patients, and 33 benign patients.

Altered cellular metabolism is a hallmark of carcinogenesis. However, current standard-of-care detection for HSILs lacks the incorporation of metabolic biomarkers. Our imaging approach relies on NAD(P)H and FP signals, providing critical information about tissue metabolism. HSILs exhibit distinct metabolic features, including a loss of spatial heterogeneity in tissue oxido-reductive state and a lower redox potential (Fig. 1D-G, Fig. 2F). These findings align with previous studies using engineered epithelial tissues and ex vivo tissue biopsies, reinforcing the robustness of our observations^17,28^.

To explore in more depth the metabolic landscape of HSILs, we conducted gene set enrichment analysis relying on publicly available epithelial-cell-enriched microarray datasets. The results indicate the occupation of a complex metabolic profile that includes overexpression of OXPHOS, glycolysis, and fatty acid metabolism along with higher levels of cellular stress (Fig. 5)^34,36,38^. Previous work by our group aimed to characterize the effects of several metabolic processes on 2P optical readouts^32^. Our observations of a decreased redox potential and an increase in mitochondrial clustering for HSIL tissues are consistent with enhanced glycolysis and fatty acid oxidation, since both are associated with decreased redox ratio values and enhanced mitochondrial clustering, i.e. more fragmented mitochondria. In contrast, enhancements in oxidative phosphorylation lead to higher redox ratio and lower mitochondrial clustering levels.

Enhanced levels of glycolysis lead to lower levels of utilization of NADH in the mitochondria and enhanced levels of NADH in the cytosol, both of which contribute to a lower redox ratio. Fragmented mitochondria are also typically more prevalent when they are not utilized aggressively for energy production. The NADH produced during fatty acid oxidation can either bind directly to the electron transport chain or be shuttled to the cytoplasm, replenishing cytosolic NADH, which is depleted in a rapidly proliferating cell^41,42^. Activation of Myc, mTOR, and HIF pathways, which are all overexpressed in HSIL tissues (Fig. 5), are tightly associated with cellular growth and proliferation; they also serve as activators for many glycolytic genes and glucose transporters^38,43–45^. Observations in cellular anabolism are also consistent with the known mechanisms of HPV viral oncoproteins E6 and E7. Oncoprotein E6 degrades p53, a key regulator of the Myc, mTOR, and HIF-1α pathways^46,47^. Oncoprotein E7 has been shown to dimerize pyruvate kinase type M2, increasing the rate of cellular proliferation and nucleotide synthesis^48^. The fact that the overall optical redox ratio of HSILs is reduced while the mitochondrial clustering is enhanced relative to non-HSILs indicates that while HSILs may produce more energy via oxidative phosphorylation to meet ATP demands, they break down glucose and fatty acids at even higher rates to maintain high levels of molecular biosynthesis. Higher levels of NADPH biosynthesis that are expected to occur as cells attempt to mitigate higher levels of oxidative stress (as indicated by UPR and antioxidant defense pathway upregulation) are also consistent with lower optical redox ratios. A greater concentration of cytosolic NAD(P)H is consistent with the significant decrease in NAD(P)H intensity in HSIL tissues, since the quantum efficiency of unbound NAD(P)H is two-to ten-fold lower than that of bound NAD(P)H, typically prevalent in mitochondria^14^ (Supplemental Figure S8). Furthermore, a decrease in NAD(P)H intensity is consistent with increased NAD(P)H consumption that would accompany an increase in gene expression for ETC Complex I subunits (Supplemental Figure S8, S9A). As for the decrease in FP intensity, underexpression of electron transfer flavoprotein dehydrogenase (ETFDH) directly prevents the oxidation of ETF, resulting in an increased proportion of non-fluorescent FADH_2_^49^ (Supplemental Figure S8, S9B). Underexpression of the GDP/ADP-forming subunit alpha of succinate-CoA ligase (SUCLG1) indirectly decreases FP intensity, by inhibiting the generation of succinate, the substrate for ETC Complex II, which utilizes FADH_2_ as a reducing equivalent^50^ (Supplemental Figure S8, S9B). Thus, the integration of optical and genomic results allows improved understanding of overall metabolic function. Nevertheless, we note that such functional insights are provided with micron scale resolution by label-free, two-photon imaging, yielding additional important information regarding the loss of spatial metabolic heterogeneity across the depth of the HSIL epithelia.

Current standard of care histopathological diagnosis relies on expert interpretation of tissue morphology, only after visual inspection of the cervix using non-specific contract agents, which results in a false positive biopsy rate of up to 94%^10^. Not only does 2PM allow for the non-invasive surveillance of multiple tissue regions, which has been shown to improve HSIL sensitivity up to 35%^7^, but overall, our study suggests that classification utilizing 2P morphofunctional biomarkers may achieve higher diagnostic accuracy (Fig. 3A)^8^. Specifically, we achieved a high 90.8 ± 6.1% sensitivity and 72.3 ± 11.3% specificity of HSIL detection by integrating metabolic and morphological 2P-derived metrics from finely sampled, full-thickness epithelia. Importantly, even when discriminating HSILs from immature non-HSIL cases, sensitivity and specificity were preserved (Fig. 3B). When using only 2 measurements from a superficial optical section, such as 12-µm, and a more basal optical section, such as 72-µm, we achieved a high sensitivity (91.4 ± 12.0%) and specificity (77.5 ± 12.6%) of detection, demonstrating the potential for rapid, sub-micrometer resolution screening and laying the groundwork for future clinical applications.

Despite the novel findings, the present study poses several limitations. The main limitation of the study is that conclusions regarding *in vivo* diagnosis are drawn from freshly excised human cervical tissue biopsies. The data acquired here remains clinically relevant due to the extensive efforts that have been made to preserve the *in vivo* condition, such as regularly hydrating the sample and limiting imaging time. By limiting the imaging time to maintain clinical relevance, we suffer in the quality of our autofluorescence signals and the range of locations which we can sample. NAD(P)H and FP autofluorescence is inherently weak. The use of 6-frame averaged images allows us to collect data of suitable SNR from several regions of interest. The use of deep-learning-based denoising algorithms would have the potential to improve the SNR of our images without the need to integrate multiple frames^51^.

The findings provided in this study motivate several areas of future investigation. The use of deep-learning-based classification methods that can integrate quantitative 2P image data and qualitative information, such as patient HPV-type, age, menstrual status, and menopausal status, have the potential to improve diagnosis^52^. Future predictive models that utilize 2P image data can also leverage morphological and metabolic measurements from the tissue stroma. ECM remodeling and patient immune response are dictated by the complex interactions that occur between epithelial cells and stromal cells^20,53–55^. 2PM can capture high-resolution, spatially preserved signatures of stromal autofluorescence and collagen SHG in 3D, in situ. Characterizing the tissue stroma and immune response may provide insights into why some lesions progress into invasive carcinomas. In fact, a quantitative multiphoton melanoma index, which integrates depth-dependent variations in epithelial autofluorescence and collagen SHG intensity, have been used to characterize skin cancer lesions imaged *in vivo*^56,57^. We have further shown that the lack of depth dependent mitochondrial clustering variations and N:C variations as assessed from analysis of in vivo NAD(P)H 2P images can differentiate human melanoma and basal cell carcinoma lesions from healthy skin^26^. Spectroscopic studies have highlighted the presence of similar epithelial and stromal autofluorescence changes associated with oral, esophageal, lung, and colorectal cancers. Portable, multi-modal, 2PM systems for assessment of morphofunctional characteristics of excised tissues at the bedside have been reported already for breast cancer^58^, lung cancer^59^, and head and neck squamous cell carcinoma^60^.

In summary, this study demonstrates the potential for rapid and robust cervical HSIL detection using non-destructive, high-resolution 2P measurements. We reveal that a loss in metabolic and morphological heterogeneity is a fundamental indicator of high-grade precancerous changes, even when comparisons are made with metaplastic and low-grade precancerous tissues. We highlight that such morphofunctional homogeneity can be captured when 2P images acquired at only two distinct epithelial depths, indicating the potential for acquiring the needed information rapidly. Using GSEA, we demonstrate that 2PM images capture functional shifts towards a more complex metabolic state that involves enhanced glycolysis, FAO, and OXPHOS. Overall, this study establishes the potential to translate non-destructive, depth-resolved, high-resolution 2P imaging to improve detection of human cervical HSILs through the quantitative assessment of spatially resolved cellular metabolic function and morphology metrics.

## Data and Code Availability

All data and codes are available from the lead contact upon reasonable request.

## Acknowledgments

We acknowledge support from the National Institute of Biomedical Imaging and Bioengineering (R01 EB030061), the National Institute of Health, Office of the Director (S10 OD021624), and the National Cancer Institute for funding this work (R03 CA235053). We would like to thank the following medical providers for their support in patient recruitment and biopsy acquisition: Alison Vogell, MD, Danielle Roncari, MD, MPH, Jenny Ruan, MD, Chenchen Sun, MD, Megan Evans, MD, MPH, Laura Baecher-Lind, MD, MPH, and Jennie Mastroianni, NP. We thank Adriana Sánchez-Hernandez for the coregistration of imaging and histology locations. We thank Paula Josephs for maintaining and compiling the patient demographic information. We also thank the Tufts Medical Center Biorepository, and specifically Karla Murga, for supporting biopsy transportation logistics. Finally, and importantly, we would like to thank the patients who consented to participate in this study.

## Author Contribution

I.G. conceived and designed the study. H.-T.T. and A.L.Z. coordinated patient recruitment and supported biopsy acquisition. Under the guidance of I.G., C.M.P. performed the imaging experiments, imaging data analysis, and imaging statistical analysis. F.R.D. consulted on imaging statistical analyses. E.M.G. and N.J. rendered all histopathological diagnoses. Under the guidance of A.P. and A.B., P.S. performed metabolic pathway analysis and the relevant statistical analyses. C.M.P. and P.S. prepared figures. C.M.P compiled and drafted the manuscript with assistance from P.S., I.G., and A.B. All authors have reviewed and approved the manuscript.

