## Supplemental Materials for "Rapid, high-resolution, non-destructive assessments of metabolic and morphological homogeneity uniquely identify high-grade cervical precancerous lesions"

* Corresponding Author

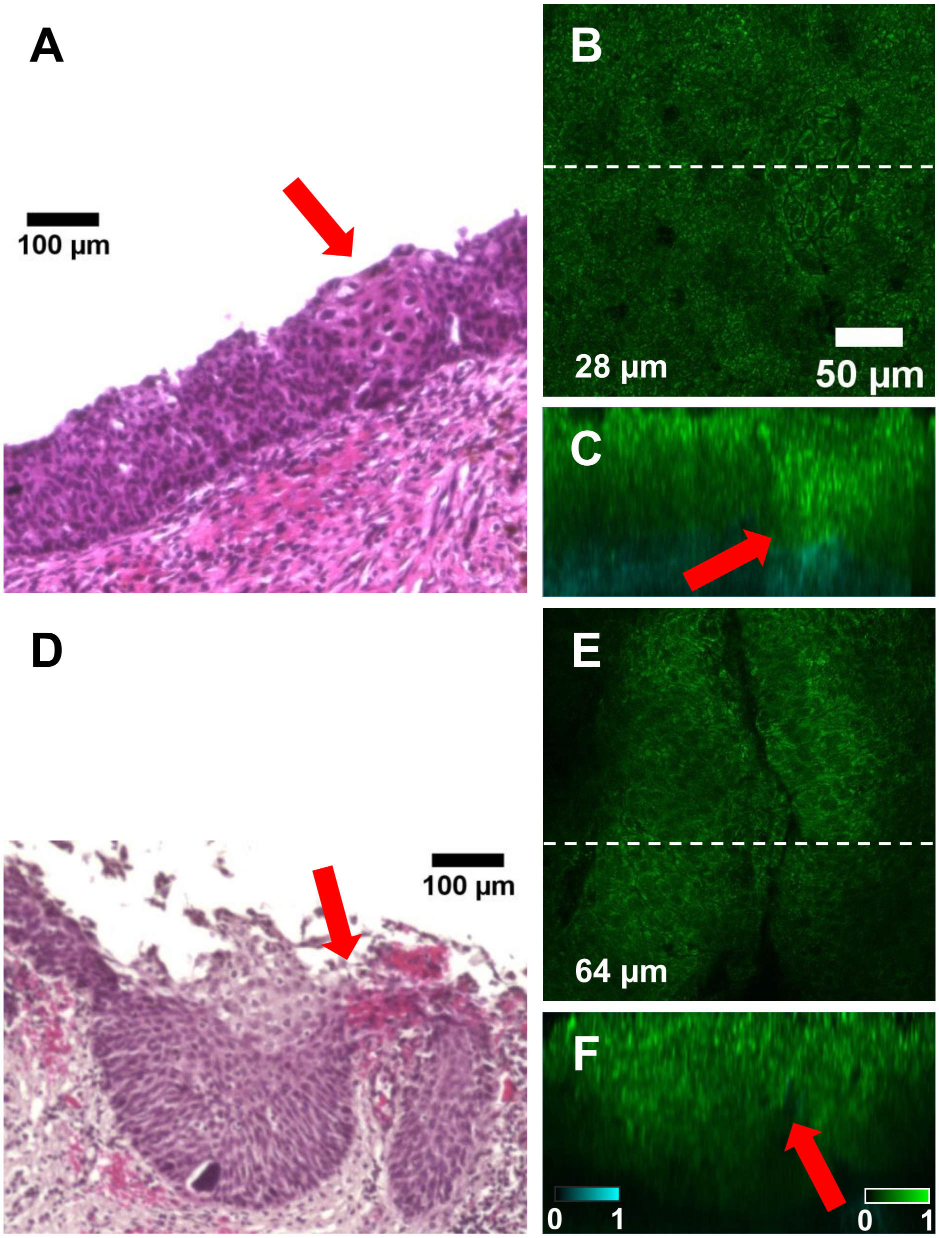


**Figure S1. Image and histology coregistration.** *En face* TPEF image locations (B, E) and hematoxylin and eosin-stained tissue cross-sections (A, D) were coregistered using methods described previously[1] and briefly in Supplemental Methods. One-to-one coregistration was conducted for 2 benign, 1 LSIL, and 2 HSIL tissue biopsies. Highlighted here is coregistration from the 2 HSIL tissues biopsies. TPEF cross sections (C, F) were rendered using Fiji (National Institute of Health) Volume Viewer and taken from the locations specified by the white dotted lines in B and E, respectively. NAD(P)H and collagen SHG were pseudo-colored green and cyan, respectively. Red arrows point to coregistered features of interest.

**Table S1. Patient exclusion criteria.**

| Characteristic | | Cohort N = 26 |
|---|---|---|
| Quality Control | Excessive Tissue Oxidation | 5 (19.2%) |
| | Stripped Epithelium | 3 (11.5%) |
| | Endocervix Only | 5 (19.2%) |
| | Mucous Only | 4 (15.4%) |
| | Testing | 1 (3.9%) |
| Specimen Access | Communication Breakdown | 6 (23.1%) |
| | Cancer Evidence | 2 (7.7%) |

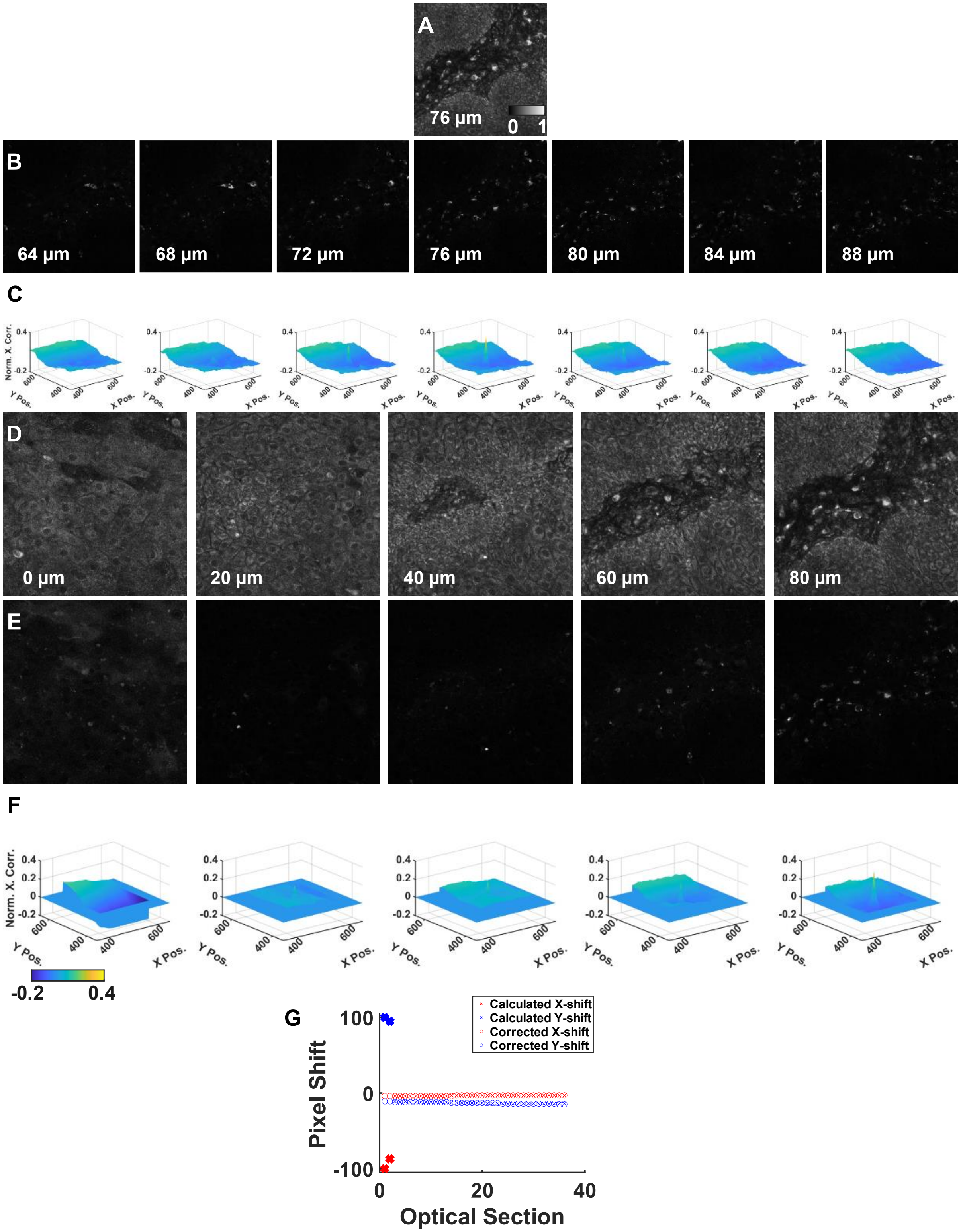
A
76 µm
0 1
B
64 µm
68 µm
72 µm
76 µm
80 µm
84 µm
88 µm
C
Norm. X. Corr.
Y Pos.
X Pos.
D
0 µm
20 µm
40 µm
60 µm
80 µm
E
F
-0.2
0.4
G
Calculated X-shift
Calculated Y-shift
Corrected X-shift
Corrected Y-shift
Pixel Shift
Optical Section

**Figure S2. Image coregistration.** A reference optical section from the 755/775 nm excitation/ 525 nm emission channel was selected for the coregistration of 755/775 and 860 nm excitation images along the depth-axis. A reference image (A) was selected based on the presence of hyperfluorescent cell signatures or cytokeratin autofluorescence, signatures that appear consistently between excitation channels. The reference image was compared with the corresponding 860 nm excitation/ 525 nm emission channel images from the same depth, three previous depths, and three following depths (B). The image pair that generated the largest normalized 2D cross-correlation value, as calculated by the MATLAB normxcorr2 built-in function, determined whether a depth shift was present between excitation channels (C). No depth shift is present for the highlighted example. After co-registering image channels in depth, normalized 2D cross-correlation maps were generated between all image pairs from the 755/775 nm excitation/ 525 nm emission (D) and 860 nm excitation/525 nm emission channels (E). The location of the normalized 2D cross-correlation peak determined how to translate the 860 nm excitation images for coregistration. Normalized 2D cross-correlation maps were limited to the central 200 x 200 pixels to exclude edge artifacts (F). The translation shift for a particular optical section was maintained if the value was within three times the difference of the median shift and minimum shift for all optical sections. In the present example, the calculated shifts for optical sections one and two are replaced for linearly interpolated values (G).

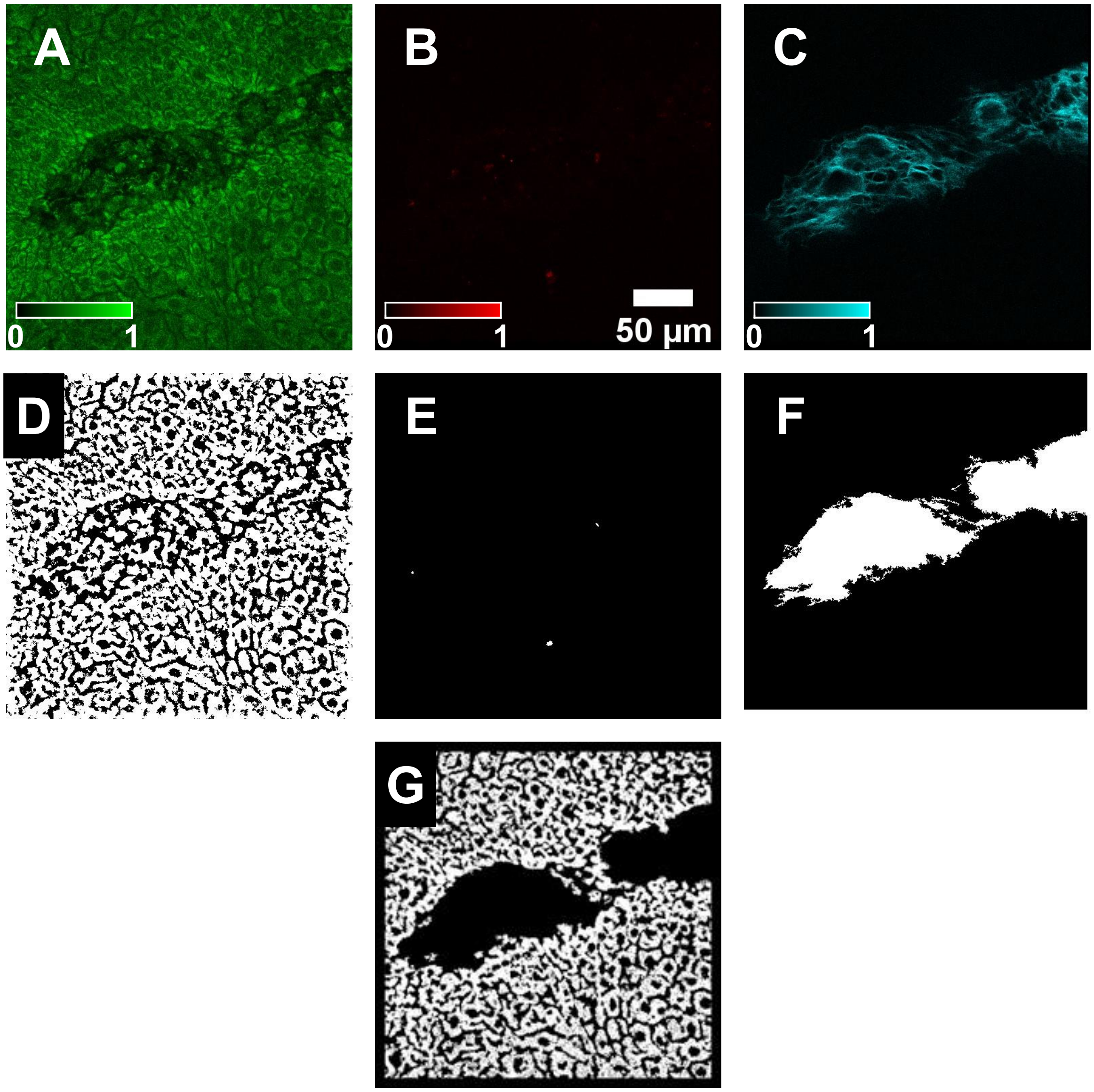


**Figure S3. Masking.** Image acquisition channels attributed to NAD(P)H (A), hyperfluorescent dead or dying cells (B), and collagen SHG (C) were masked to isolate cellular cytoplasm. Features corresponding to nuclei and interstitial spaces were removed through frequency filtering and Otsu-based intensity thresholding (D). Signals attributed to dead and dying cells were enhanced and removed using a low-pass filter and stack-level Otsu thresholding (E). Collagen-positive pixel regions were identified using a stack-level Otsu's global threshold; Holes in collagen-positive regions were filled and once a pixel was identified as collagen-positive it was masked for the remainder of the image volume (F). Cytoplasm-positive pixels (G) were identified by combining (D) with the inverse of (E) and (F) and removing a 15-pixel border around the image.

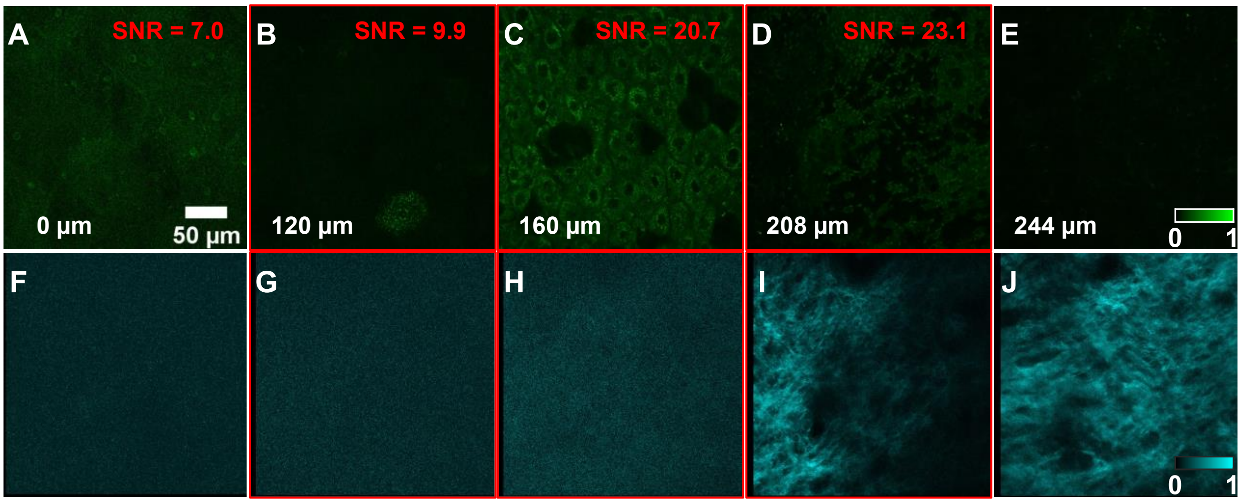


**Figure S4. Optical section inclusion criteria.** Signal-to-noise ratio (SNR) was calculated from NAD(P)H images (A – E). The first optical section with an SNR greater than nine (B) was the first optical section considered for analysis. SNR thresholding filtered superficial optical sections (A) with weak autofluorescence emanating from cytokeratins, as opposed to mitochondrial NAD(P)H. The last optical section with greater than 30% cell coverage (D) was the last optical section considered for analysis. Percent cell coverage was determined by the presence of collagen SHG (F – J). The epithelium for the highlighted ROI was defined as the region between 120 to 208 µm. Intraepithelial optical sections are highlighted in red (B – D, G - I).

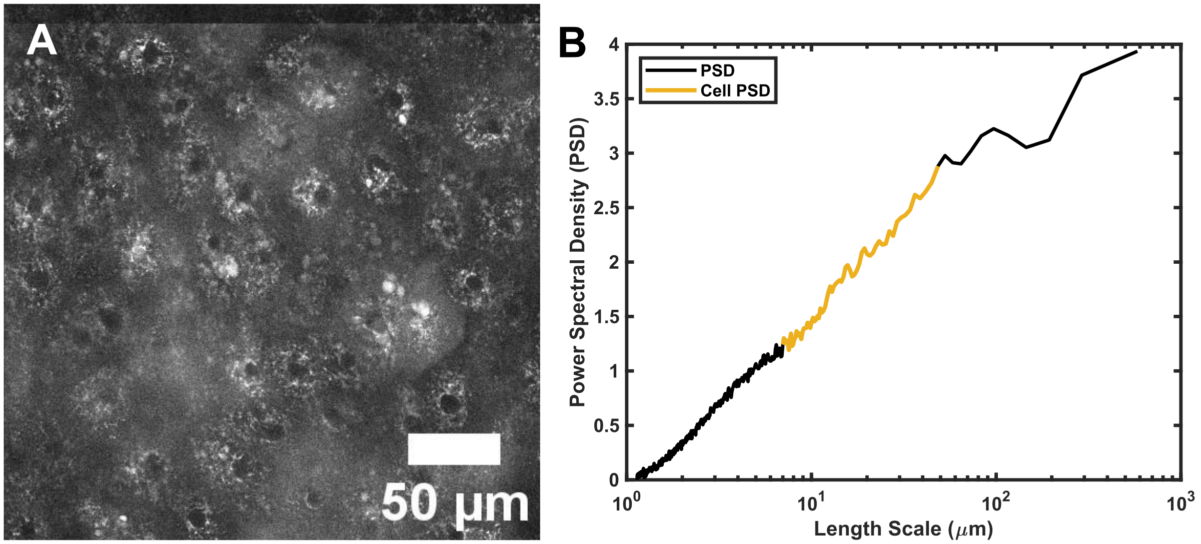


**Figure S5. Representative image power spectral density (PSD).** A representative integrated intensity image (A) and the corresponding PSD, where values for cellular length scales are highlighted in yellow (B).

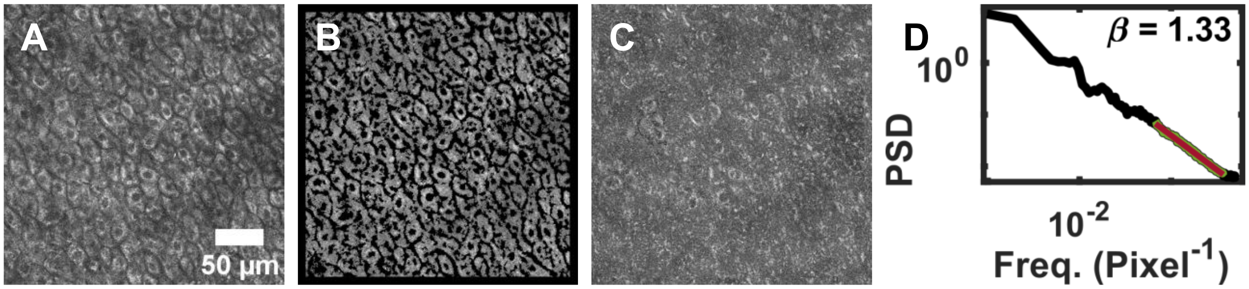


**Figure S6. Mitochondrial clustering analysis.** NAD(P)H images (A) were masked to remove autofluorescence signatures emanating from cellular nuclei and tissue interstitial spaces (B). Signal attributed to cellular cytoplasm was digitally object cloned to fill a full field of view without overwriting cytoplasm-positive pixels (C). The power spectral density (PSD) from 5 randomly digitally object cloned images was fit for frequencies corresponding to <8.5-µm length scales with a line of the form R(k) = Ak-β, where β is reported as a metric of mitochondrial clustering (D). In (D), the representative PSD calculated from (C) is black, the PSD for frequencies corresponding to <8.5-µm length scales is green, and the inverse power-law fit is red.

**Table S2. Effects of stepwise metric inclusion on QDA AUC for HSIL vs. mature-containing non-HSIL dataset.**

| Metric | AUC Metric Only | AUC Metric + (1) | AUC Metric + (1, 2) | AUC Metric + (1 - 3) | AUC Metric + (1 - 4) | AUC Metric + (1 - 5) | AUC Metric + (1 - 6) |
|---|---|---|---|---|---|---|---|
| Diff. Gradient (1) | 0.6988 | - | - | - | - | - | - |
| RR IQR Range (2) | 0.6977 | 0.8170 | - | - | - | - | - |
| Ep. Thickness (3) | 0.6532 | 0.7879 | 0.8508 | - | - | - | - |
| Median RR (4) | 0.6232 | 0.7263 | 0.8302 | 0.8828 | | - | - |
| Median β (5) | 0.5669 | 0.6979 | 0.8206 | 0.8690 | 0.8951 | - | - |
| β Variability (6) | 0.5469 | 0.7263 | 0.8164 | 0.8591 | 0.8848 | 0.9024 | - |
| RR Variability (7) | 0.5324 | 0.7325 | 0.8197 | 0.871 | 0.8897 | 0.9138 | 0.9235 |

**Table S3. Effects of stepwise metric inclusion on QDA AUC for HSIL vs. immature non-HSIL dataset.**

| Metric | AUC Metric Only | AUC Metric + (1) | AUC Metric + (1, 2) | AUC Metric + (1 - 3) | AUC Metric + (1 - 4) | AUC Metric + (1 - 5) | AUC Metric + (1 - 6) |
|---|---|---|---|---|---|---|---|
| RR IQR Range (1) | 0.7116 | - | - | - | - | - | - |
| Diff. Gradient (2) | 0.6819 | 0.8062 | - | - | - | - | - |
| Ep. Thickness (3) | 0.5830 | 0.7284 | 0.8320 | - | - | - | - |
| Median RR (4) | 0.5648 | 0.7425 | 0.8234 | 0.8587 | | - | - |
| Median β (5) | 0.5419 | 0.7233 | 0.8086 | 0.8558 | 0.8757 | - | - |
| RR Variability (6) | 0.5203 | 0.7059 | 0.8071 | 0.8505 | 0.8732 | 0.9001 | - |
| β Variability (7) | 0.4832 | 0.7218 | 0.8071 | 0.8322 | 0.8574 | 0.8796 | 0.9072 |

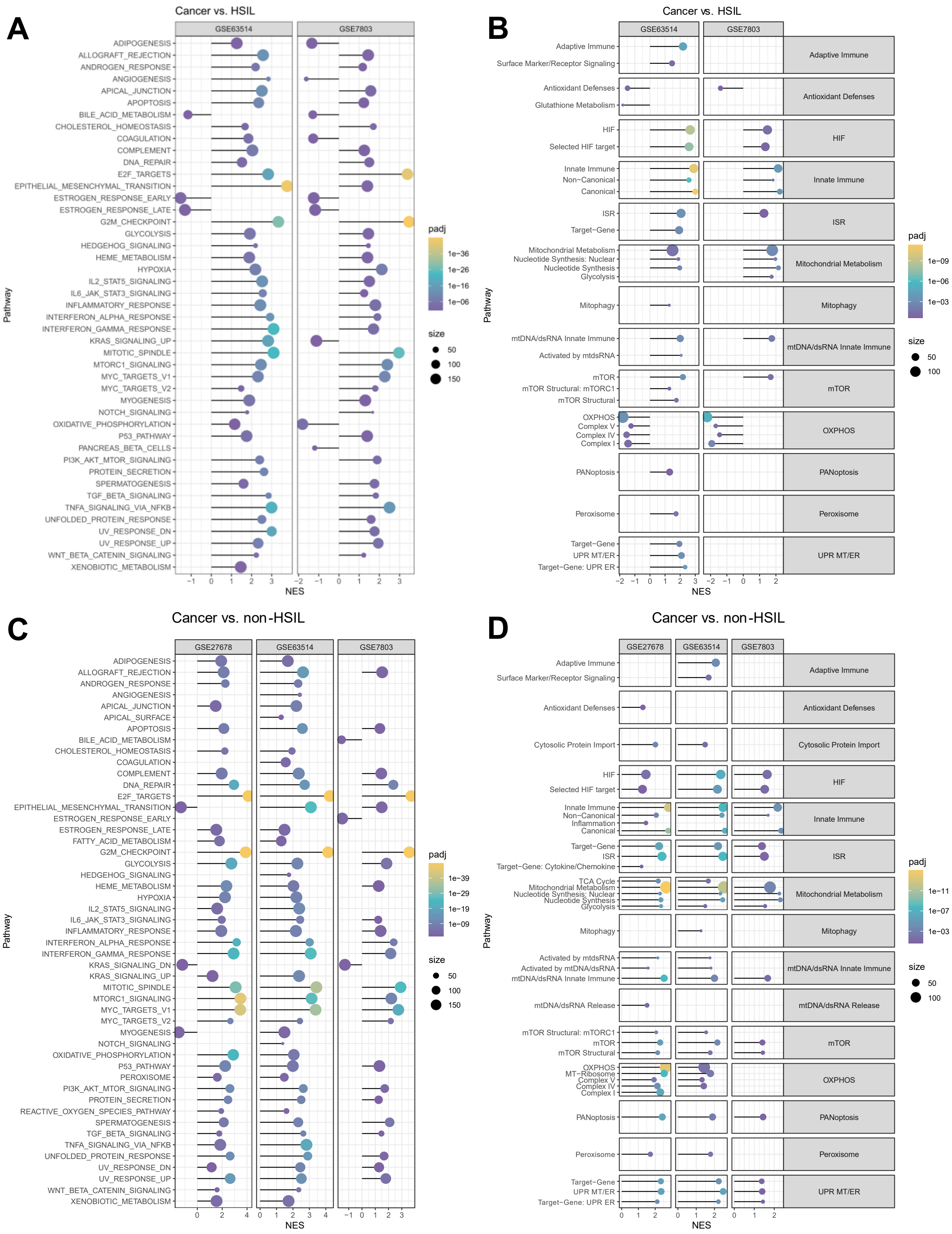

171

172

**Figure S7: Differential pathway expression in invasive cervical cancer.** Fast gene set enrichment analysis (FGSEA) using Hallmark (A, C) and custom mitochondrial (B, D) gene sets was used to compare cancerous tissue with HSIL (A – B) and non-HSIL (C – D) tissues in three independent datasets (GSE27678, GSE63514, and GSE7803). A positive normalized enrichment score (NES) indicates that a particular pathway is upregulated in cancerous tissues. Dot size corresponds to the size of the corresponding gene set. The dot pseudocolor is scaled based on the calculated Benjamini-Hochberg (BH)-adjusted p-value.

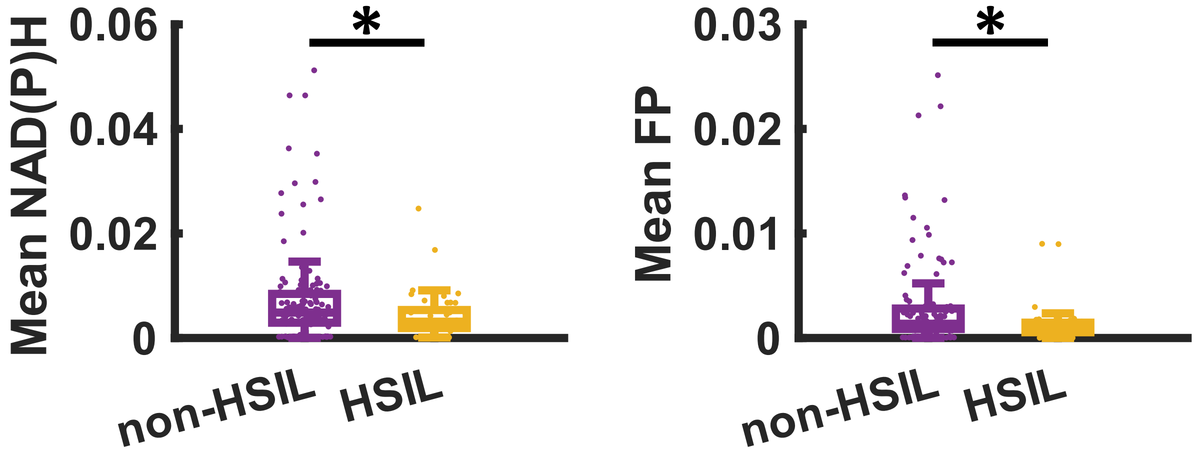


**Figure S8. NAD(P)H and FP Intensity.** HSILs tissues were characterized by a significantly lower NAD(P)H and FP intensity.

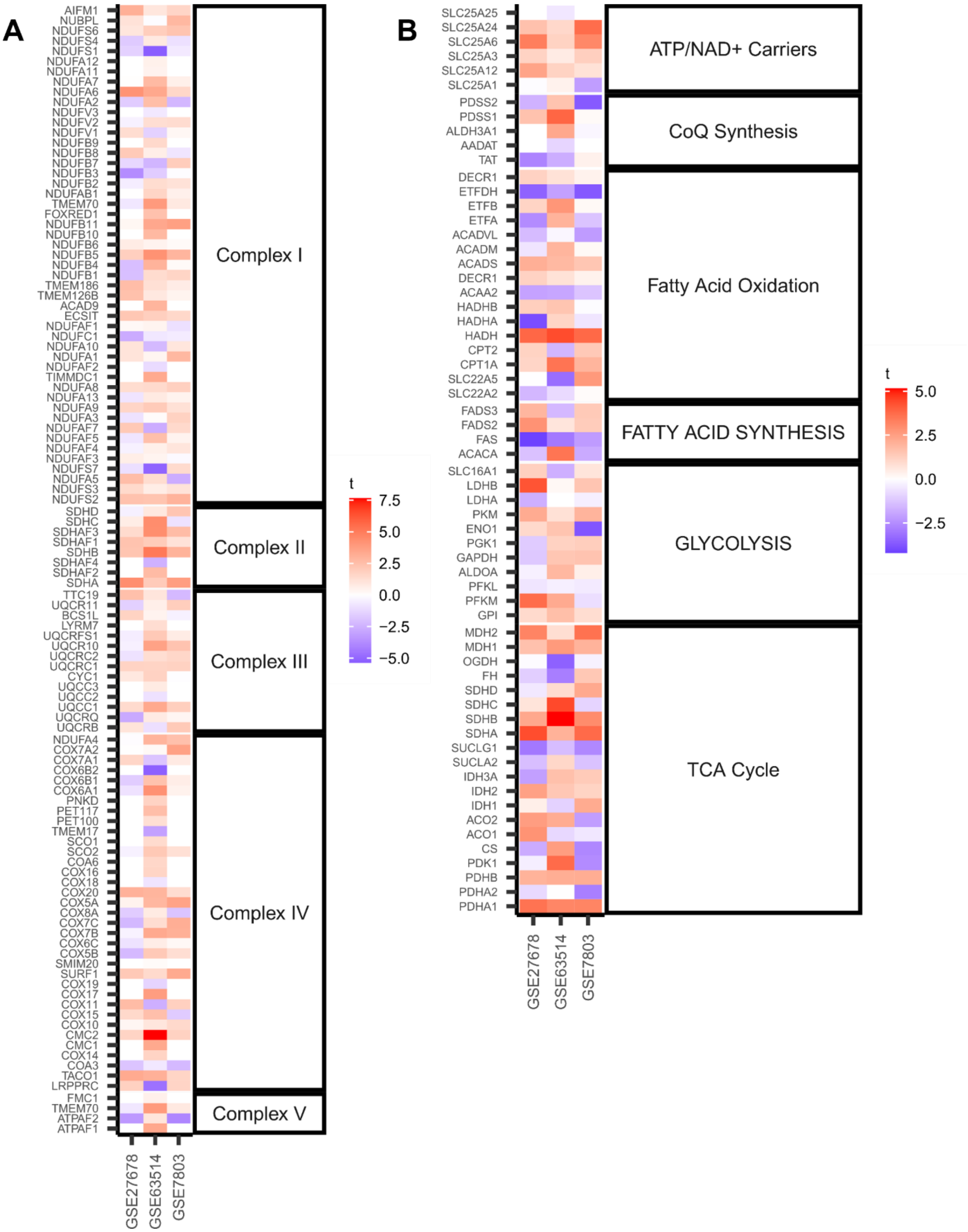
A
AIFM1
NUBPL
NDUFS6
NDUFS4
NDUFS1
NDUFA12
NDUFA11
NDUFA7
NDUFA6
NDUFA2
NDUFV3
NDUFV2
NDUFV1
NDUFB9
NDUFB8
NDUFB7
NDUFB3
NDUFB2
NDUFAB1
TMEM70
FOXRED1
NDUFB11
NDUFB10
NDUFB6
NDUFB5
NDUFB4
NDUFB1
TMEM186
TMEM126B
ACAD9
ECSIT
NDUFAF1
NDUFC1
NDUFA10
NDUFA1
NDUFAF2
TIMMDC1
NDUFA8
NDUFA13
NDUFA9
NDUFA3
NDUFAF7
NDUFAF5
NDUFAF4
NDUFAF3
NDUFS7
NDUFA5
NDUFS3
NDUFS2
SDHD
SDHC
SDHAF3
SDHAF1
SDHB
SDHAF4
SDHAF2
SDHA
TTC19
UQCR11
BCS1L
LYRM7
UQCRFS1
UQCR10
UQCRC2
UQCRC1
CYC1
UQCC3
UQCC2
UQCC1
UQCRQ
UQCRB
NDUFA4
COX7A2
COX7A1
COX6B2
COX6B1
COX6A1
PNKD
PET117
PET100
TMEM17
SCO1
SCO2
COA6
COX16
COX18
COX20
COX5A
COX8A
COX7C
COX7B
COX6C
COX5B
SMIM20
SURF1
COX19
COX17
COX11
COX15
COX10
CMC2
CMC1
COX14
COA3
TACO1
LRPPRC
FMC1
TMEM70
ATPAF2
ATPAF1
Complex I
Complex II
Complex III
Complex IV
Complex V
GSE27678
GSE63514
GSE7803
t
7.5
5.0
2.5
0.0
−2.5
−5.0
B
SLC25A25
SLC25A24
SLC25A6
SLC25A3
SLC25A12
SLC25A1
PDSS2
PDSS1
ALDH3A1
AADAT
TAT
DECR1
ETFDH
ETFB
ETFA
ACADVL
ACADM
ACADS
DECR1
ACAA2
HADHB
HADHA
HADH
CPT2
CPT1A
SLC22A5
SLC22A2
FADS3
FADS2
FAS
ACACA
SLC16A1
LDHB
LDHA
PKM
ENO1
PGK1
GAPDH
ALDOA
PFKL
PFKM
GPI
MDH2
MDH1
OGDH
FH
SDHD
SDHC
SDHB
SDHA
SUCLG1
SUCLA2
IDH3A
IDH2
IDH1
ACO2
ACO1
CS
PDK1
PDHB
PDHA2
PDHA1
ATP/NAD+ Carriers
CoQ Synthesis
Fatty Acid Oxidation
FATTY ACID SYNTHESIS
GLYCOLYSIS
TCA Cycle
GSE27678
GSE63514
GSE7803
t
5.0
2.5
0.0
−2.5

**Figure S9. Differential gene expression heatmaps.** Differential gene expression was assessed between HSIL and non-HSIL tissues in three independent datasets (GSE27678, GSE63514, and GSE7803). A positive t-score indicates that a particular gene is overexpressed in HSIL tissues compared to non-HSIL tissues. Genes belonging to the custom mitochondrial oxidative phosphorylation (A) and mitochondrial metabolism (B) pathways are illustrated.

### Supplemental Methods:

#### *Histology and Image Coregistration*

The coregistration of imaging locations and locations from which hematoxylin and eosin-stained tissue cross sections were taken allowed for a 1:1 histopathological diagnosis to be rendered. For the five biopsies in which there was histology and image coregistration, the periphery of the tissue surface was marked with tattoo inks prior to imaging. Specifically, three markings were made with a tattoo needle on the imaging surface with red, green, and blue tattoo inks (Dynamic Color). Tattoo ink locations were imaged using reflected light generated by a 488 nm diode laser. Following image acquisition, tattoo ink and imaging locations were mapped using positional data saved in the microscope software. Accounting for ~20% shrinkage during tissue fixation and paraffin-embedding, imaging locations relative to ink positions could be approximated for sectioning.

#### *Patient Exclusion Criteria*

There were eight occasions in which no tissue was taken from the hospital. Two hysterectomy specimens presented evidence of cancer and therefore no samples were taken for imaging. Breakdowns in communication, such as the use of Lugol's iodine for colposcopic inspection, uterine specimen morcellation, or a lack of immediate notification from operating room staff following hysterectomy specimen removal, can be attributed to six other occasions where tissue was not taken for imaging. There were 18 occasions in which tissues were taken for imaging, but data was not taken or excluded. One specimen was used to optimize imaging parameters. Five specimens exhibited considerable oxidation in which little-to-no NAD(P)H autofluorescence was present in the epithelia. Three specimens had stripped epithelium, as evidenced by the presence of collagen on the tissue surface. Five specimens only contained endocervix, as was evidenced by single cell layer optical images, and confirmed via histopathological diagnosis. Four specimens were either too small or did not contain intact tissue from which the tissue epithelium could be located (Supplemental Table S1).

*Image Channel Coregistration*

Although 755/775- and 860-nm excitation occurred in rapid succession, small deformations could still be observed in the tissue. For this reason, 3D spatial coregistration was necessary to accurately assess tissue metabolic state at each image pixel. A qualitatively similar optical section between the 755/775- and 860-nm excitation, 525/50 nm emission channels was specified (Supplemental Figure S2A). The reference optical section in the 755/775 nm excitation channel was compared with seven, 860 nm excitation images, the three optical sections before, the three optical sections after, and the optical section at the same reference depth (Supplemental Figure S2B). The 860 nm excitation optical section with the highest normalized cross-correlation determined whether a depth shift was present between excitation channels (Supplemental Figure S2C). Normalized 2D cross-correlation was calculated between images using the built-in MATLAB normxcorr2 function. Normalized cross-correlation analysis yields a quantitative measure of similarity, with values ranging from -1 to 1, for all potential translational shifts of two images. After co-registering for depth, normalized cross-correlation analysis was conducted for each paired image within the stack (Supplemental Figure S2D-E). The 2D shift in the 860 nm excitation image was determined by the spatial location of maximal normalized cross-correlation (Supplemental Figure S2F). If the total shift in x- and y- for a particular optical section was greater than three-times the difference of the median shift and minimum shift of all optical sections, the shift was omitted for a linearly interpolated value. For the example illustrated in Supplemental Figure S2, linearly interpolated values were leveraged to shift optical sections one and two (Supplemental Figure S2G). An erroneous shift in optical section one can be attributed to a lack of bright features between the 755/775- and 860-nm images, as visualized in the $1^{st}$ panels (0-µm) of Supplemental Figure S2D-E. Such dissimilarity results in a normalized 2D cross-correlation peak at the window edge instead of a central location (Supplemental Figure S2F).

*Masking*

Signals attributed to cellular nuclei and interstitial tissue regions were removed using NAD(P)H images. NAD(P)H images were subject to contrast-limited adaptive histogram equalization prior to the serial application of two Gaussian and one 3rd order Butterworth bandpass filter. Nuclei and interstitial regions were removed by using the lower level of a two-level Otsu’s global threshold for binarization. Hyperfluorescent cell signals were isolated by applying the higher level of a two-level Otsu’s global threshold to Gaussian low-pass filtered images from the 860 nm excitation, 624/40 nm emission channel. A mask was generated from collagen SHG images to delineate the epithelium and remove stromal autofluorescence contributions. Collagen-positive pixels were identified by applying the lower level of a two-level Otsu’s global threshold to SHG images. Gaps within collagen-positive pixels were filled, and once a pixel was identified as collagen-positive, it was masked for the remainder of the stack. A low, empirically determined noise threshold of 0.0002 was also implemented to aid in the identification of collagen for ROIs in which the stroma was not extensively sampled. The inverted hyperfluorescent cell mask and inverted collagen SHG mask were combined with the NAD(P)H mask to define cytoplasmic regions. Features in a 15-pixel band around the edge of each image were masked to remove frequency filtering artifacts created during nuclear and interstitial feature removal. Small, less than 25-pixel objects were removed to finalize the mask, omitting residual fragments.

*Comparing mean NAD(P)H and mean FP*

Mean NAD(P)H and mean FP values were extracted for cytoplasm-positive pixels from each intraepithelial optical section. Nested t-tests were used to make statistical comparisons for the mean NAD(P)H and mean FP of each ROI as a function of depth. Both mean NAD(P)H and mean FP intensity were significantly lower in HSIL tissues (Supplemental Figure S8).

*Gene set enrichment analysis*

The three datasets accessioned from GEO were further investigated to evaluate metabolic comparisons between invasive cancer, and HSIL and non-HSIL groups. The datasets corresponding to GSE27678, GSE 63514, and GSE7803 contained 29, 28, and 21 cancerous tissue samples, respectively. Comparisons between cancer and HSIL tissues could not be made for GSE27678 due to the use of two different microarray platforms during data acquisition. Normalized enrichment scores and statistical significance of custom mitochondrial and Hallmark gene sets were evaluated using fGSEA as previously described. Glycolysis was upregulated in cancer vs non-HSIL tissues in all three datasets for both Hallmark and custom mitochondrial gene sets.

Signaling pathways implicated in the enhanced activation of glycolytic pathways such as mammalian target of rapamycin (mTOR), Myc, and hypoxia inducible factor (HIF) were also upregulated in cancerous tissues. Genes sets implicated in oxidative catabolic processes such as the tricarboxylic acid (TCA) cycle, fatty acid oxidation, and oxidative phosphorylation (OXPHOS) were upregulated in 2 of the 3 datasets. As for tissue anabolism, gene sets implicated in nucleotide synthesis and cellular proliferation (E2F targets, G2M checkpoint) were upregulated in all cancerous tissue datasets (Supplemental Figure S7C – D).

For cancerous vs. HSIL tissues, Hallmark gene sets indicated enhanced glycolytic activity in cancerous tissues for both datasets. Relevant glycolytic Hallmark signaling pathways (mTOR, Myc, Hypoxia/HIF) were also significantly upregulated. Using the custom mitochondrial pathways, glycolysis and mTOR gene sets were upregulated in only one of the datasets. Custom mitochondrial OXPHOS gene sets were downregulated in cancer tissues compared to HSIL tissues. This trend was preserved in one of the datasets, while reversed in the other, when using the Hallmark gene sets. Gene sets pertaining to cellular anabolism such as Hallmark E2F targets, Hallmark G2M checkpoint, and custom mitochondrial nucleotide synthesis were upregulated in cancer tissues (Supplemental Figure S7A – B).
